# Thirty five novel nsSNPs may effect on *ADAMTS13* protein leading to Thrombotic thrombocytopenic purpura (TTP) using bioinformatics approach

**DOI:** 10.1101/759787

**Authors:** Tebyan A. Abdelhameed, Arwa I. Ahmed, Mujahed I. Mustafa, Amel N. Eltayeb, Fatima A. Abdelrhman, Amal B. Ahmed, Najla B. Ahmed, Hiba Y. Khadir, Adla M. Hassan, Huda K. Mohamed, Soada A. Osman, Mustafa Elhag, Mohamed A. Hassan

**Author notes:** Authors contribution is equally (Tebyan A. Abdelhameed, Arwa I. Ahmed). Corresponding author: Tebyan A. Abdelhameed.

## Abstract

**Background:** Genetic polymorphisms in the *ADAMTS13* gene are associated with thrombotic thrombocytopenic purpura or TTP, a life-threatening microangiopathic disorder. This study aims to predict the possible pathogenic SNPs of this gene and their impact on the protein structure and function using insilico methods.

**Methods:** SNPs retrieved from the NCBI database were analyzed using several bioinformatics tools. The different algorithms applied collectively to predictthe effect of single nucleotide substitution on both structure and function of the *ADAMTS13* protein.

**Results:** Fifty one mutations were found to be highly damaging to the structure and function of the protein. Of those, thirty five were novel nsSNPs not previously reported in the literature.

**Conclusion:** According to our analysis we found thirty five nsSNPs effects on *ADAMTS13* protein leading to thrombotic thrombocytopenic purpura using computational approach. Bioinformatics tools are vital in prediction analysis, making use of increasingly voluminous biomedical data thereby providing markers for screening or for genetic mapping studies.

## INTRODUCTION

Thrombotic thrombocytopenic purpura (TTP), also known as Upshaw – Schulman syndrome (USS) is one of microangiopathic disorder which is a life threating thrombotic microangiopathy [1–4], caused by widespread of microthrombi composed from highly reactive high molecular von Willebrand factor (vWF) and platelets due to deficiency of the vWF-cleaving metalloprotease encoded by *ADAMTS13* gene [1, 5, 6]. *ADAMTS13* (a disintegrin and metalloprotease with thrombospondin type 1 repeats, member 13), is highly specific multi-domain plasma reprolysin-like metalloprotease [4, 7–9], located on chromosome 9 (9q34) [10], which is normally synthesized in the kidney (major side of production), liver and endothelium cells [11, 12]. The disease is characterized by thrombocytopenia, hemolytic anemia, neurologic signs, fever, organ ischemia and subsequently impaired function of different organs especially the kidneys and the central nervous system (CNS) [4, 5, 11, 13–16].

The congenital TTP is usually occurring due to mutations in *ADAMTS13,* while the acquired occur due to autoantibodies against *ADAMTS13* [11–13]. Several studies revealed that the single nucleotide polymorphisms may be associated with reduced *ADAMTS13* secretion and activity, and some mutations in *ADAMTS13* gene responsible from phenotype expression of TTP characteristics [2, 7, 10–13, 17–22]. Although there are some researchers studied the mutations in *ADAMTS13* [10, 12, 14, 16, 20, 21, 23] but the genetic mechanism of some mutations remain unclear. Our study aimed to define the impact of various SNPs in *ADAMTS13gene* using a bioinformatics approach and assessing the effect of these SNPs on the structure and function of *ADAMTS13* protein that may be implicated in the disease susceptibility. Recently, bioinformatics approach make huge differences in science through providing massive information about genes, proteins, domains, SNPs, and many more. It is valuable to know genes functions and their interaction with other genes so the pathways in normal and abnormal case, genetic evolution to know who is closest to whom, proteins function so the structure and proprieties and SNPs and its association with disorder which all poured in health line. This study is regarded as the first computational study to predict the effects of SNPs on *ADAMTS13* protein function and structure.

## METHODS

### Data mining

The data for human *ADAMTS13* gene were collected from the National Center for Biological Information (NCBI) website [24]. The SNP information (SNP ID) of the *ADAMTS13* gene was retrieved from the NCBI dbSNP (http://www.ncbi.nlm.nih.gov/snp/) and the protein ID and its sequence was collected from Swiss-Prot databases with the accession number: (Q76LX8). (http://expasy.org/) [25].

### SIFT

SIFT (Sorting Intolerant from Tolerant) is an online computational tool [26] to detect a harmful non-synonymous single-base nucleotide polymorphism (nsSNP). SIFT predicts whether an amino acid replacement affects protein function. SIFT prediction is based on the grade of conservation of amino acid residues in sequence alignments resulting from closely related sequences, collected through PSI-BLAST. SIFT can be applied to naturally occurring non-synonymous polymorphisms or laboratory induced missense mutations, the genetic mutation that causes a single amino acid substitution (AAS) in a protein sequence subsequently changing the carrier’s phenotype and health status. SIFT expects whether these substitutions affect protein function by using sequence homology, SIFT predicts the effects of all possible substitutions at each position in the protein sequence. Furthermore, the algorithm performs a comprehensive search in protein repositories to find the tolerance of each candidate compared to the conserved counterparts. Non-synonymous reference SNPs identity (nsSNPs ID) were downloaded from online dbSNPs of NCBI and then submitted to SIFT. Results are expressed as damaging (deleterious) or benign (Tolerated) depending on cutoff value 0.05; as values below or equal to (0.0-0.04) predicted to be damaging or intolerant while (0.051) is benign or tolerated, then the damaging SNPs were reanalyzed by Polyphen software which predicts the effect of mutations on both structural and functional sides. It is available at (http://sift.bii.a-star.edu.sg/).

### Polyphen-2

Polyphen-2 is a software tool [27] used to predict the possible impact of an amino acid substitution on both structure and function of a human protein by analysis of multiple sequence alignment and protein 3D structure, in addition, it calculates position-specific independent count scores (PSIC) for each of two variants, and then calculates the (PSIC) scores difference between two variants. The higher a PSIC score difference, the higher the functional impact a particular amino acid substitution is expected to have. Prediction outcomes could be categorized as probably damaging, possibly damaging or benign according to the value of PSIC as it ranges from (0_1); values closer to zero considered benign while values closer to 1 considered probably damaging and also it can be indicated by a vertical black marker inside a color gradient bar, where green is benign and red is damaging, nsSNPs that predicted to be intolerant by Sift has been submitted to Polyphen as protein sequence in FASTA format that obtained from UniproktB /Expasy after submitting the relevant ensemble protein (ESNP) there, and then we entered position of mutation, native amino acid and the new substituent for both structural and functional predictions. PolyPhen version 2.2.2 is available at the website. (http://genetics.bwh.harvard.edu/pph2/index.shtml)

### Provean

Provean is a software tool which predicts whether an amino acid substitution or indel has an impact on the biological function of a protein. It is useful for purifying sequence variants to identify nonsynonymous or indel variants that are predicted to be functionally important [28]. Variants with a score equal to or below −2.5 are considered “deleterious” while variants with a score above −2.5 are considered “neutral.” It is available at (https://rostlab.org/services/snap2web/).

### SNAP2

Functional effects of mutations are predicted with SNAP2 [29]. SNAP2 is a trained classifier that is based on a machine learning device called “neural network”. It distinguishes between effect and neutral variants/non-synonymous SNPs by taking a variety of sequence and variant features into account. The most important input signal for the prediction is the evolutionary information taken from an automatically generated multiple sequence alignment. Also, structural features such as predicted secondary structure and solvent accessibility are considered. If available also annotation (i.e. known functional residues, pattern, regions) of the sequence or close homologs are pulled in. In a cross-validation over 100,000 experimentally annotated variants, SNAP2 reached sustained two-state accuracy (effect/neutral) of 82% (at an AUC of 0.9). In our hands, this constitutes an important and significant improvement over other methods. It is available at (https://rostlab.org/services/snap2web/).

### PHD-SNP

PHD-SNP is an online Support Vector Machine (SVM) based classifier optimized to predict if a given single point protein mutation can be classified as disease-related or as a neutral polymorphism [30]. It is available at: (http://http://snps.biofold.org/phd-snp/phdsnp.html)

### SNPs&Go

SNPs&Go is an algorithm developed in the Laboratory of Biocomputing at the University of Bologna directed by Prof. Rita Casadio. SNPs&Go is an accurate method that, starting from a protein sequence, can predict whether a variation is disease related or not by exploiting the corresponding protein functional annotation. SNPs&Go collects in unique framework information derived from protein sequence, evolutionary information, and function as encoded in the Gene Ontology terms, and outperforms other available predictive methods [30]. It is available at (http://snps.biofold.org/snps-and-go/snps-and-go.html)

### Pmut

PMUT a web-based tool for the annotation of pathological variants on proteins, allows the fast and accurate prediction (approximately 80% success rate in humans) of the pathological character of single point amino acidic mutations based on the use of neural networks [31]. It is available at (http://mmb.irbbarcelona.org/PMut).

### I-Mutant 3.0

I-Mutant 3.0 Is a neural network based tool for the routine analysis of protein stability and alterations by taking into account the single-site mutations [32]. The FASTA sequence of protein retrieved from UniProt is used as an input to predict the mutational effect on protein stability. It is available at (http://gpcr2.biocomp.unibo.it/cgi/predictors/I-Mutant3.0/I-Mutant3.0.cgi).

### Project Hope

Project hope (version 1.0 is an online webserver used to search protein 3D structures by collecting structural information from a series of sources, including calculations on the 3D protein structure (if available), sequence annotations in UniproktB and predictions from DAS servers. Protein sequences were submitted to project hope server in order to analyze the structural and conformational variations that have resulted from single amino acid substitution corresponding to single nucleotide substitution. It is available at (http://www.cmbi.ru.nl/hope) [33].

### UCSF (University of California at San Francisco) Chimera

UCSF Chimera (https://www.cgl.ucsf.edu/chimera/) is a highly extensible program for interactive visualization and analysis of molecular structures and related data, including density maps, supramolecular assemblies, sequence alignments, docking results, trajectories, and conformational ensembles. High-quality images and animations can be generated. Chimera includes complete documentation and several tutorials. Chimera is developed by the Resource for Biocomputing, Visualization, and Informatics (RBVI), supported in part by the National Institutes of Health (P41-GM103311) [34].

### GeneMANIA

We submitted genes and selected from a list of data sets that they wish to query. GeneMANIA approach to know protein function prediction integrate multiple genomics and proteomics data sources to make inferences about the function of unknown proteins [35]. It is available at (http://www.genemania.org/).

## RESULTS

### *ADAMTS13* gene investigation using dbSNP/NCBI

This study identified the total number of nsSNP in Homo sapiens located in the coding region of *ADAMTS13* gene, were investigate in dbSNP/NCBI Database [24]. Out of 1291 SNPs there are 821 nsSNPs (missense mutations).

### Functional analysis of *ADAMTS13 gene* using SIFT, Polyphen-2, Provean, SNAP2, PHD-SNP, SNP&GO, PHD-SNP and P-Mut softwares

821 nsSNPs (missense mutations) were submitted to SIFT server, PolyPhen-2 server, Provean sever and SNAP2 respectively, 323 SNPs were predicted to be deleterious in SIFT server. In PolyPhen-2 server, our result showed that 491 were found to be damaging (133 possibly damaging and 358 probably damaging showed deleterious). In Provean server our result showed that 393 SNPs were predicted to be deleterious. While in SNAP2 server our result showed that 418 SNPs were predicted to be Effect. The differences in prediction capabilities refer to the fact that every prediction algorithm uses different sets of sequences and alignments. In table (2) we have submitted four positive results which are 66 results from SIFT, PolyPhen-2, Provean and SNAP2 to observe the disease causing one by SNP&GO, PHD-SNP and P-Mut servers.

**Table (1):** Damaging or Deleterious or effect nsSNPs associated variations predicted by various softwares:

| Amino Acid Change | SIFT |  | Polyphen |  | PROVEAN |  | SNAP2 |  |
| --- | --- | --- | --- | --- | --- | --- | --- | --- |
|  | Prediction | Score | Prediction | Score | Score | Prediction | Prediction | Score |
| S207R | D | 0 | probably damaging | 1 | -4.694 | Deleterious | Effect | 72 |
| C208S | D | 0 | probably damaging | 1 | -9.654 | Deleterious | Effect | 73 |
| G215S | D | 0 | probably damaging | 1 | -5.792 | Deleterious | Effect | 39 |
| H234Q | D | 0 | probably damaging | 1 | -7.412 | Deleterious | Effect | 64 |
| D235Y | D | 0 | probably damaging | 1 | -8.077 | Deleterious | Effect | 74 |
| S263Y | D | 0 | probably damaging | 1 | -5.39 | Deleterious | Effect | 56 |
| P301R | D | 0 | probably damaging | 1 | -7.772 | Deleterious | Effect | 45 |
| G302A | D | 0 | probably damaging | 1 | -5.447 | Deleterious | Effect | 34 |
| C311Y | D | 0 | probably damaging | 1 | -10.612 | Deleterious | Effect | 73 |
| L335P | D | 0 | probably damaging | 1 | -6.907 | Deleterious | Effect | 69 |
| P353L | D | 0 | probably damaging | 1 | -9.201 | Deleterious | Effect | 43 |
| W390C | D | 0 | probably damaging | 1 | -12.291 | Deleterious | effect | 65 |
| R398C | D | 0 | probably damaging | 1 | -7.608 | Deleterious | effect | 74 |
| R398H | D | 0 | probably damaging | 1 | -4.753 | Deleterious | effect | 97 |
| G403V | D | 0 | probably damaging | 1 | -8.572 | Deleterious | effect | 44 |
| R409W | D | 0 | probably damaging | 1 | -7.608 | Deleterious | effect | 82 |
| C411Y | D | 0 | probably damaging | 1 | -10.457 | Deleterious | effect | 70 |
| P416H | D | 0 | probably damaging | 1 | -8.281 | Deleterious | effect | 63 |
| G419W | D | 0 | probably damaging | 1 | -7.569 | Deleterious | effect | 79 |
| G419A | D | 0 | probably damaging | 1 | -5.662 | Deleterious | effect | 55 |
| G419V | D | 0 | probably damaging | 1 | -8.506 | Deleterious | effect | 59 |
| R421C | D | 0 | probably damaging | 1 | -5.298 | Deleterious | effect | 46 |
| C423R | D | 0 | probably damaging | 1 | -11.441 | Deleterious | effect | 57 |
| G425D | D | 0 | probably damaging | 1 | -6.674 | Deleterious | effect | 54 |
| G425V | D | 0 | probably damaging | 1 | -8.581 | Deleterious | effect | 48 |
| C450R | D | 0 | probably damaging | 1 | -11.856 | Deleterious | effect | 78 |
| C483Y | D | 0 | probably damaging | 1 | -10.57 | Deleterious | effect | 53 |
| D504N | D | 0 | probably damaging | 1 | -4.927 | Deleterious | effect | 47 |
| T506N | D | 0 | probably damaging | 1 | -4.94 | Deleterious | effect | 50 |
| C508Y | D | 0 | probably damaging | 1 | -9.405 | Deleterious | effect | 54 |
| G525D | D | 0 | probably damaging | 1 | -6.89 | Deleterious | effect | 85 |
| R660W | D | 0 | probably damaging | 1 | -4.945 | Deleterious | effect | 53 |
| S696W | D | 0 | probably damaging | 1 | -5.135 | Deleterious | effect | 50 |
| C699R | D | 0 | probably damaging | 1 | -10.987 | Deleterious | effect | 90 |
| C758W | D | 0 | probably damaging | 1 | -8.165 | Deleterious | effect | 82 |
| G759R | D | 0 | probably damaging | 1 | -5.59 | Deleterious | effect | 67 |
| G759W | D | 0 | probably damaging | 1 | -5.693 | Deleterious | effect | 47 |
| G761S | D | 0 | probably damaging | 1 | -4.454 | Deleterious | effect | 44 |
| C864R | D | 0 | probably damaging | 1 | -8.616 | Deleterious | effect | 58 |
| C908Y | D | 0 | probably damaging | 1 | -10.074 | Deleterious | effect | 82 |
| C908S | D | 0 | probably damaging | 1 | -9.16 | Deleterious | effect | 77 |
| G911V | D | 0 | probably damaging | 1 | -8.252 | Deleterious | effect | 60 |
| C951G | D | 0 | probably damaging | 1 | -10.071 | Deleterious | effect | 95 |
| R979W | D | 0 | probably damaging | 1 | -4.359 | Deleterious | effect | 82 |
| C1024G | D | 0 | probably damaging | 1 | -11.055 | Deleterious | effect | 97 |
| C1028R | D | 0 | probably damaging | 1 | -11.021 | Deleterious | effect | 85 |
| C1028Y | D | 0 | probably damaging | 1 | -10.092 | Deleterious | effect | 85 |
| G1031R | D | 0 | probably damaging | 1 | -7.22 | Deleterious | effect | 89 |
| R1035C | D | 0 | probably damaging | 1 | -6.806 | Deleterious | effect | 80 |
| C1039Y | D | 0 | probably damaging | 1 | -9.175 | Deleterious | effect | 71 |
| C1192Y | D | 0 | probably damaging | 1 | -5.868 | Deleterious | effect | 74 |

**Table (2):** Disease effect nsSNPs associated variations predicted by various softwares:

| Amino acid change | PHD |  |  | SNP & GO |  | P-MUT |  |  |
| --- | --- | --- | --- | --- | --- | --- | --- | --- |
|  | Prediction | RI | Probability | Probability | RI | Prediction | Score | Prediction |
| S207R | Disease | 6 | 0.819 | Disease | 2 | 0.596 | 0.74 (87%) | Disease |
| C208S | Disease | 7 | 0.852 | Disease | 6 | 0.819 | 0.79 (89%) | Disease |
| G215S | Disease | 7 | 0.844 | Disease | 6 | 0.813 | 0.75 (87%) | Disease |
| H234Q | Disease | 6 | 0.819 | Disease | 7 | 0.833 | 0.79 (89%) | Disease |
| D235Y | Disease | 9 | 0.928 | Disease | 8 | 0.903 | 0.83 (90%) | Disease |
| S263Y | Disease | 3 | 0.659 | Disease | 5 | 0.743 | 0.78 (88%) | Disease |
| P301R | Disease | 5 | 0.773 | Disease | 1 | 0.552 | 0.69 (86%) | Disease |
| G302A | Disease | 4 | 0.693 | Disease | 5 | 0.761 | 0.87 (91%) | Disease |
| C311Y | Disease | 9 | 0.964 | Disease | 8 | 0.897 | 0.79 (89%) | Disease |
| L335P | Disease | 7 | 0.828 | Disease | 6 | 0.799 | 0.80 (89%) | Disease |
| P353L | Disease | 4 | 0.711 | Disease | 1 | 0.528 | 0.74 (87%) | Disease |
| W390C | Disease | 5 | 0.75 | Disease | 5 | 0.751 | 0.83 (90%) | Disease |
| R398C | Disease | 7 | 0.861 | Disease | 5 | 0.738 | 0.82 (90%) | Disease |
| R398H | Disease | 5 | 0.747 | Disease | 2 | 0.592 | 0.75 (87%) | Disease |
| G403V | Disease | 6 | 0.823 | Disease | 5 | 0.738 | 0.79 (89%) | Disease |
| R409W | Disease | 9 | 0.926 | Disease | 7 | 0.859 | 0.69 (86%) | Disease |
| C411Y | Disease | 9 | 0.945 | Disease | 7 | 0.867 | 0.79 (89%) | Disease |
| P416H | Disease | 6 | 0.8 | Disease | 1 | 0.54 | 0.83 (90%) | Disease |
| G419W | Disease | 8 | 0.896 | Disease | 6 | 0.808 | 0.79 (89%) | Disease |
| G419A | Disease | 6 | 0.78 | Disease | 2 | 0.608 | 0.56 (81%) | Disease |
| G419V | Disease | 8 | 0.885 | Disease | 5 | 0.77 | 0.73 (87%) | Disease |
| R421C | Disease | 7 | 0.854 | Disease | 6 | 0.809 | 0.72 (87%) | Disease |
| C423R | Disease | 9 | 0.928 | Disease | 8 | 0.887 | 0.79 (89%) | Disease |
| G425D | Disease | 8 | 0.923 | Disease | 7 | 0.86 | 0.74 (87%) | Disease |
| G425V | Disease | 8 | 0.919 | Disease | 7 | 0.862 | 0.79 (89%) | Disease |
| C450R | Disease | 8 | 0.894 | Disease | 7 | 0.837 | 0.79 (89%) | Disease |
| C483Y | Disease | 9 | 0.956 | Disease | 8 | 0.902 | 0.79 (89%) | Disease |
| D504N | Disease | 7 | 0.834 | Disease | 6 | 0.819 | 0.73 (87%) | Disease |
| T506N | Disease | 5 | 0.761 | Disease | 5 | 0.763 | 0.74 (87%) | Disease |
| C508Y | Disease | 8 | 0.886 | Disease | 7 | 0.85 | 0.79 (89%) | Disease |
| G525D | Disease | 8 | 0.876 | Disease | 7 | 0.838 | 0.75 (88%) | Disease |
| R660W | Disease | 5 | 0.771 | Disease | 7 | 0.842 | 0.53 (80%) | Disease |
| S696W | Disease | 6 | 0.822 | Disease | 4 | 0.704 | 0.83 (90%) | Disease |
| C699R | Disease | 9 | 0.927 | Disease | 7 | 0.875 | 0.76 (88%) | Disease |
| C758W | Disease | 8 | 0.9 | Disease | 7 | 0.863 | 0.60 (82%) | Disease |
| G759R | Disease | 5 | 0.763 | Disease | 3 | 0.657 | 0.59 (82%) | Disease |
| G759W | Disease | 7 | 0.833 | Disease | 5 | 0.748 | 0.71 (86%) | Disease |
| G761S | Disease | 6 | 0.795 | Disease | 3 | 0.628 | 0.67 (85%) | Disease |
| C864R | Disease | 7 | 0.844 | Disease | 3 | 0.632 | 0.65 (84%) | Disease |
| C908Y | Disease | 9 | 0.937 | Disease | 7 | 0.864 | 0.83 (90%) | Disease |
| C908S | Disease | 8 | 0.9 | Disease | 6 | 0.821 | 0.83 (90%) | Disease |
| G911V | Disease | 8 | 0.884 | Disease | 5 | 0.738 | 0.83 (90%) | Disease |
| C951G | Disease | 8 | 0.894 | Disease | 1 | 0.556 | 0.78 (88%) | Disease |
| R979W | Disease | 6 | 0.799 | Disease | 2 | 0.578 | 0.69 (86%) | Disease |
| C1024G | Disease | 8 | 0.884 | Disease | 5 | 0.736 | 0.85 (91%) | Disease |
| C1028R | Disease | 8 | 0.879 | Disease | 6 | 0.797 | 0.85 (91%) | Disease |
| C1028Y | Disease | 8 | 0.91 | Disease | 6 | 0.8 | 0.85 (91%) | Disease |
| G1031R | Disease | 6 | 0.779 | Disease | 3 | 0.657 | 0.76 (88%) | Disease |
| R1035C | Disease | 9 | 0.927 | Disease | 2 | 0.615 | 0.83 (90%) | Disease |
| C1039Y | Disease | 9 | 0.95 | Disease | 7 | 0.85 | 0.76 (88%) | Disease |
| C1192Y | Disease | 7 | 0.827 | Disease | 0 | 0.512 | 0.55 (81%) | Disease |

In SNP&GO, PHD-SNP and P-Mut softwares were used to predict the association of SNPs with the disease. According to SNP&GO, PHD-SNP and P-Mut (57, 66 and 57 SNPs respectively) were found to be disease related SNPs. We selected the triple disease related SNPs only in 3 softwares (51 SNPs) for further analysis by I-Mutant 3.0, Table (3).

**Table (3):** stability analysis predicted by I-Mutant version 3.0:

| Substitution | I-Mutant Prediction | I-Mutant Score | I-Mutant RI |
| --- | --- | --- | --- |
| S207R | Decrease | 0.08 | 0 |
| C208S | Decrease | -0.58 | 3 |
| G215S | Decrease | -1.11 | 8 |
| H234Q | Decrease | -0.21 | 3 |
| D235Y | Increase | -0.12 | 6 |
| S263Y | Increase | 0.11 | 4 |
| P301R | Decrease | -0.83 | 5 |
| G302A | Decrease | -0.39 | 1 |
| C311Y | Decrease | -0.32 | 3 |
| L335P | Decrease | -1.46 | 2 |
| P353L | Decrease | -0.77 | 7 |
| W390C | Decrease | -1.3 | 8 |
| R398C | Decrease | -0.99 | 3 |
| R398H | Decrease | -1.17 | 7 |
| G403V | Decrease | -0.61 | 6 |
| R409W | Decrease | -0.44 | 4 |
| C411Y | Decrease | -0.35 | 3 |
| P416H | Decrease | -1.26 | 9 |
| G419W | Decrease | -0.25 | 4 |
| G419A | Decrease | -0.61 | 6 |
| G419V | Decrease | -0.53 | 6 |
| R421C | Decrease | -0.73 | 4 |
| C423R | Increase | -0.26 | 0 |
| G425D | Decrease | -0.91 | 7 |
| G425V | Decrease | -0.38 | 5 |
| C450R | Increase | 0.02 | 2 |
| C483Y | Decrease | -0.98 | 1 |
| D504N | Decrease | -1.25 | 5 |
| T506N | Decrease | -1.01 | 3 |
| C508Y | Decrease | -0.37 | 1 |
| G525D | Decrease | -0.79 | 4 |
| R660W | Decrease | -0.26 | 4 |
| S696W | Increase | 0 | 3 |
| C699R | Decrease | -0.55 | 5 |
| C758W | Decrease | -0.19 | 3 |
| G759R | Decrease | -0.46 | 4 |
| G759W | Decrease | -0.28 | 2 |
| G761S | Decrease | -0.89 | 5 |
| C864R | Decrease | -0.13 | 0 |
| C908Y | Decrease | -0.4 | 3 |
| C908S | Decrease | -0.86 | 8 |
| G911V | Decrease | -1.7 | 1 |
| C951G | Decrease | -0.98 | 5 |
| R979W | Decrease | -0.38 | 6 |
| C1024G | Decrease | -1.39 | 9 |
| C1028R | Decrease | -0.52 | 6 |
| C1028Y | Decrease | -0.38 | 3 |
| G1031R | Decrease | -0.54 | 6 |
| R1035C | Decrease | -0.73 | 5 |
| C1039Y | Decrease | -0.11 | 3 |
| C1192Y | Decrease | -0.13 | 0 |

### Prediction of Change in Stability due to Mutation Using I-Mutant 3.0 Server

I-Mutant result revealed that the protein stability decreased which destabilize the amino acid interaction (46). While five were found to be increased the protein stability (Table 3).

**Figure (1):**
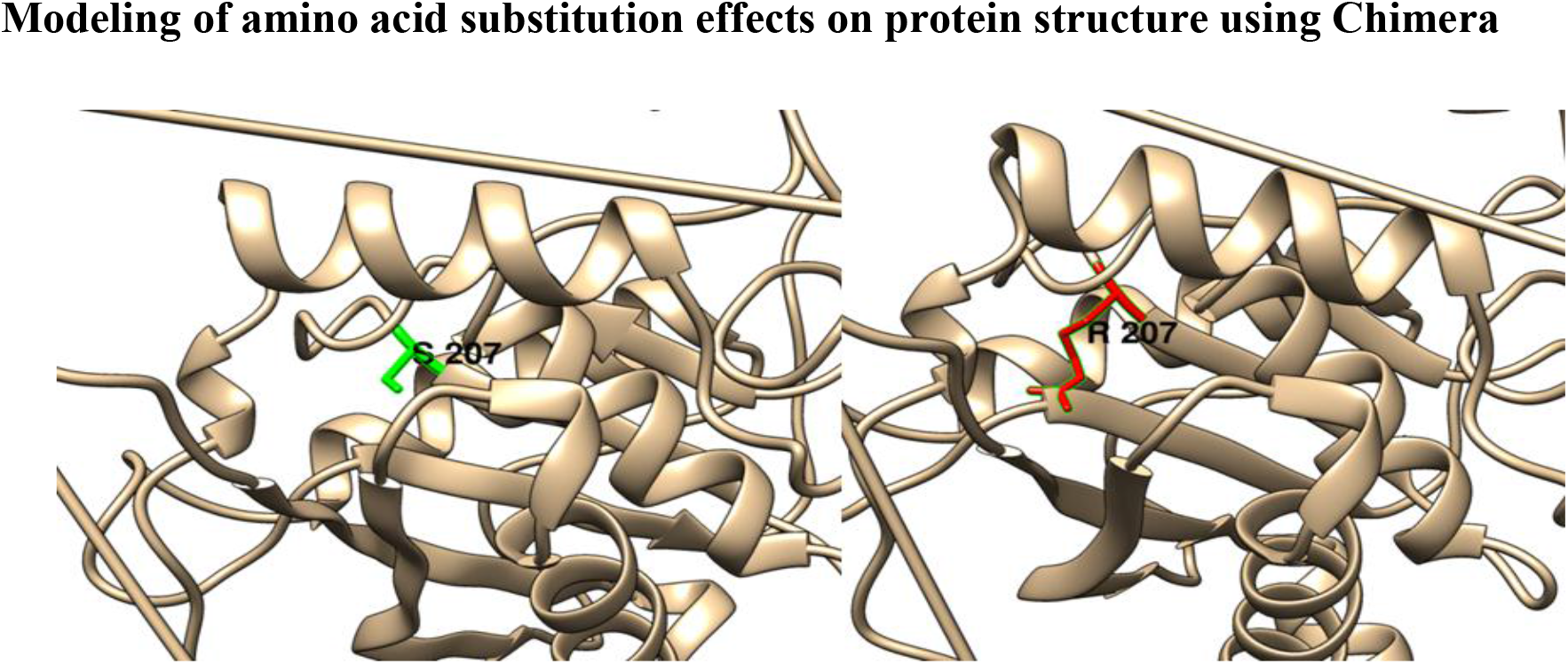
(S207R): change in the amino Serine (green box) into an Arginine (redbox) at position 207.

**Figure (2):**
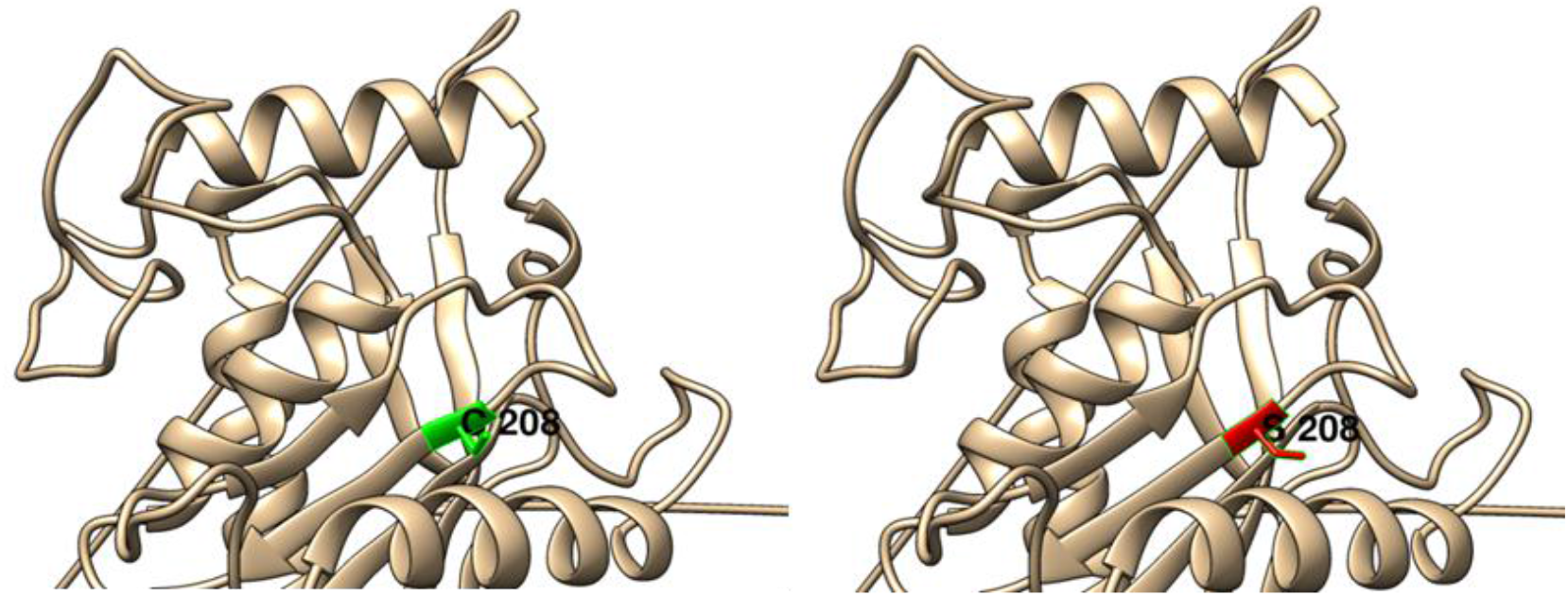
(C208S): change in the amino Cysteine (greenbox) into a Serine (red box) at position 208.

**Figure (3):**
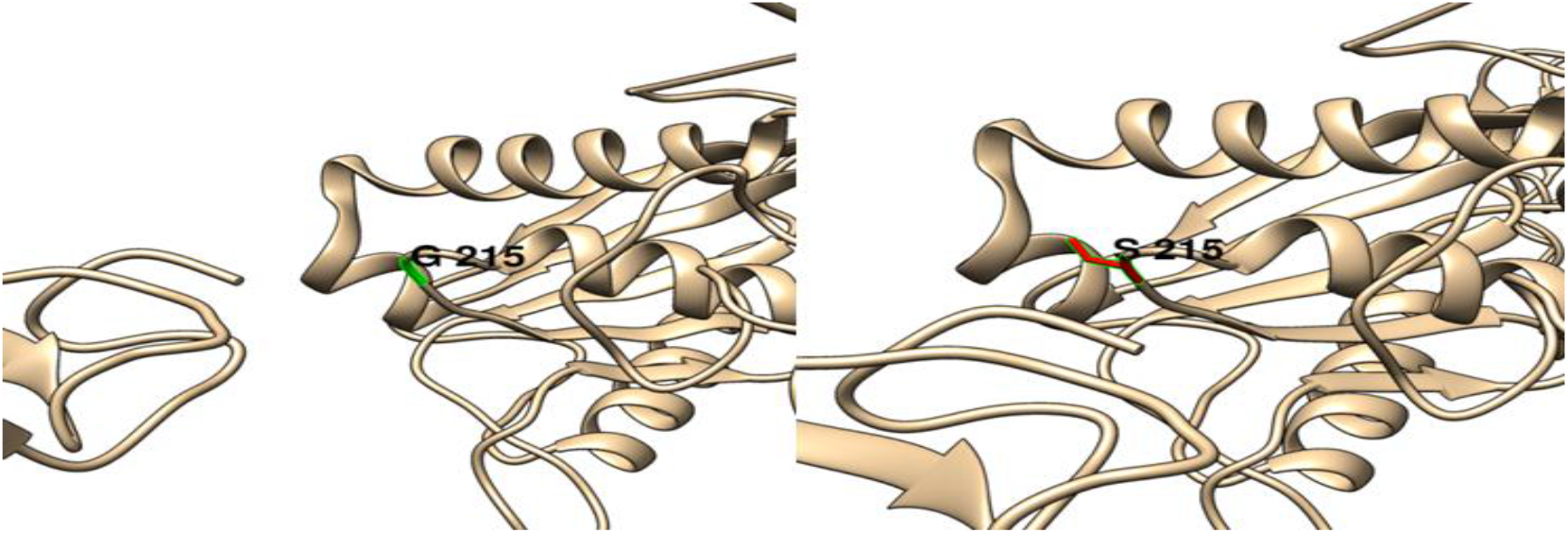
(G215S): change in the amino Glycine (green box) into a Serine (redbox) at position 215.

**Figure (4):**
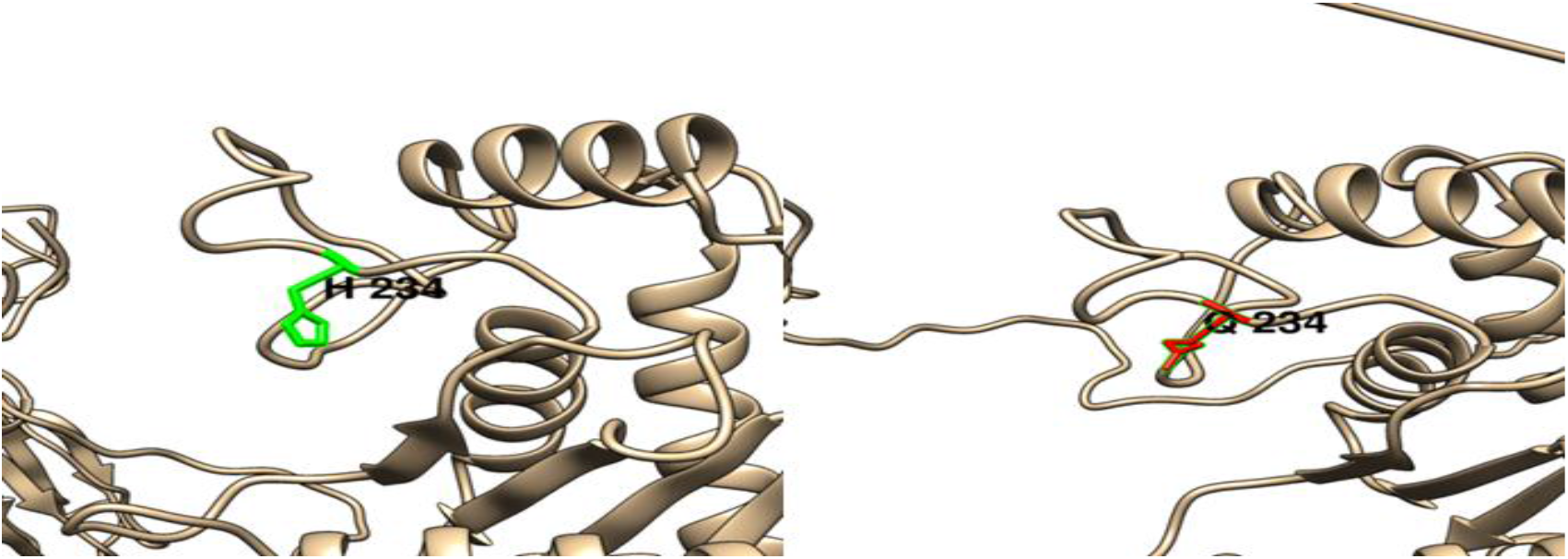
(H234Q): change in the amino Histidine (green box) into a Glutamine (red box) at position 234.

**Figure (5):**
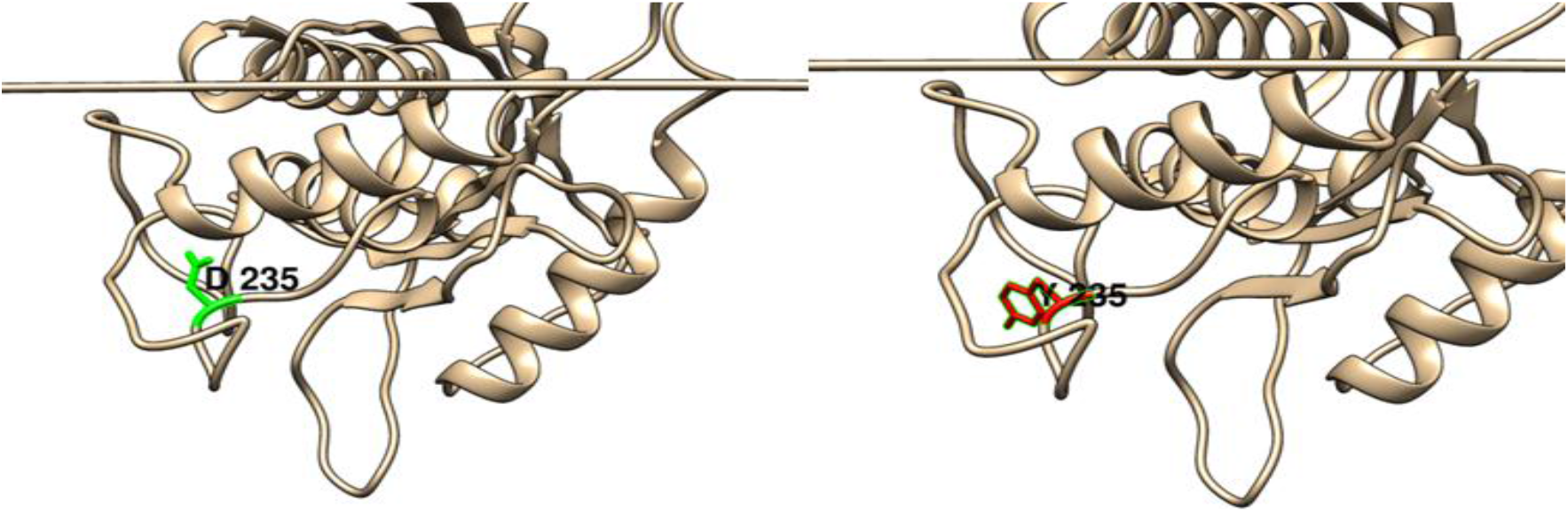
(D235Y): change in the amino Aspartic Acid (green box) into a Tyrosine (red box) at position 235.

**Figure (6):**
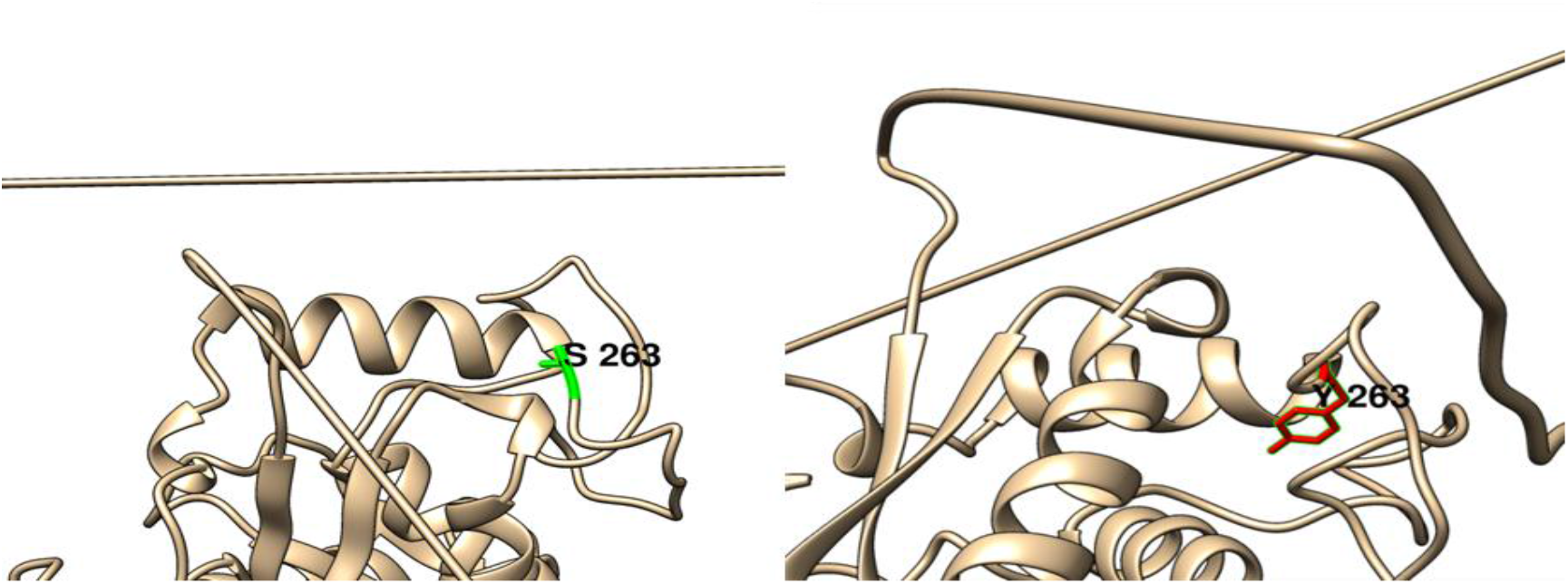
(S263Y): change in the amino Serine (green box) into a Tyrosine (red box) at position 263.

**Figure (7):**
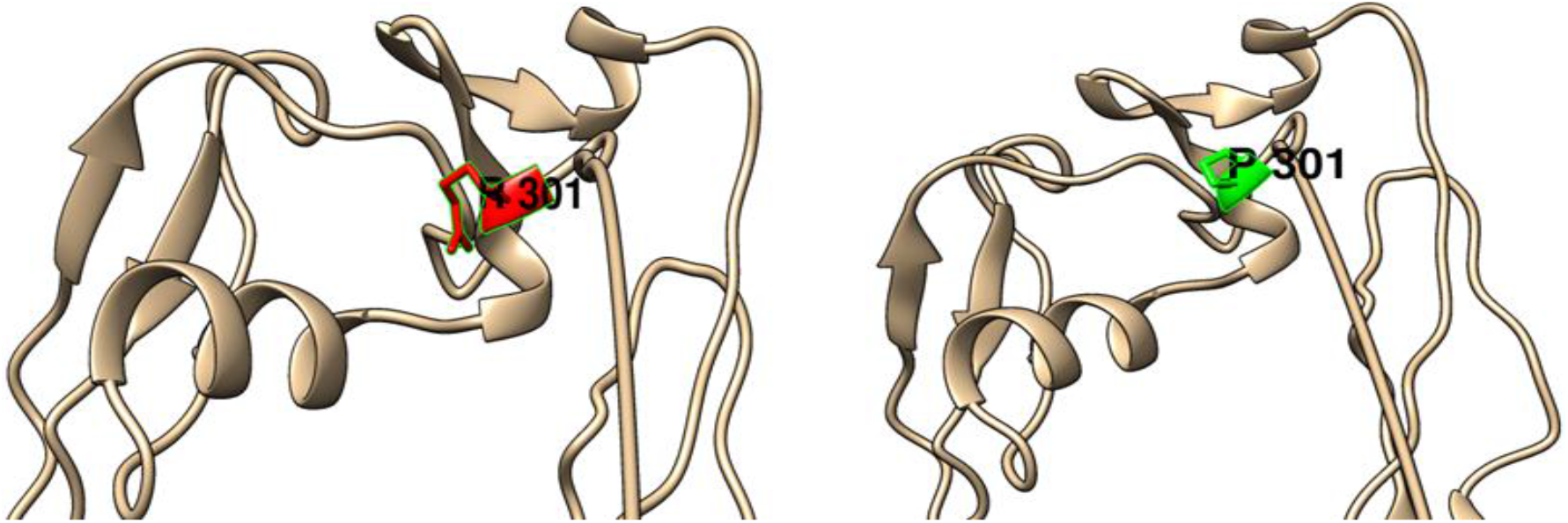
(P301R) change in the amino Proline (green box) into an Arginine (red box) at position 301.

**Figure (8):**
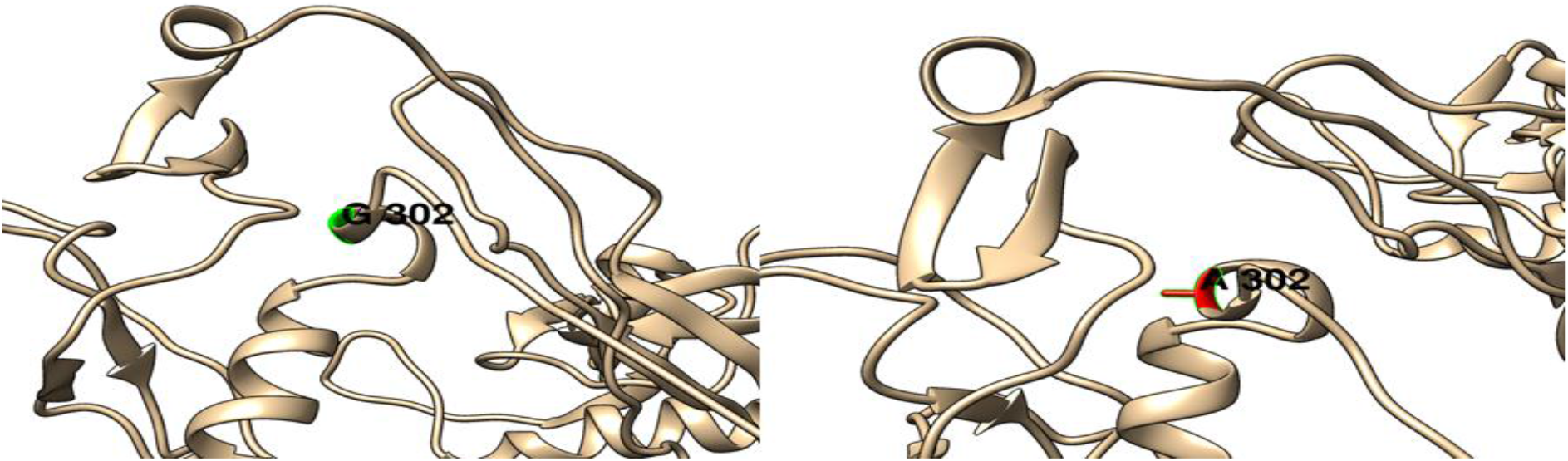
(G302A): change in the amino Glycine (green box) into an Alanine (red box) at position 302.

**Figure (9):**
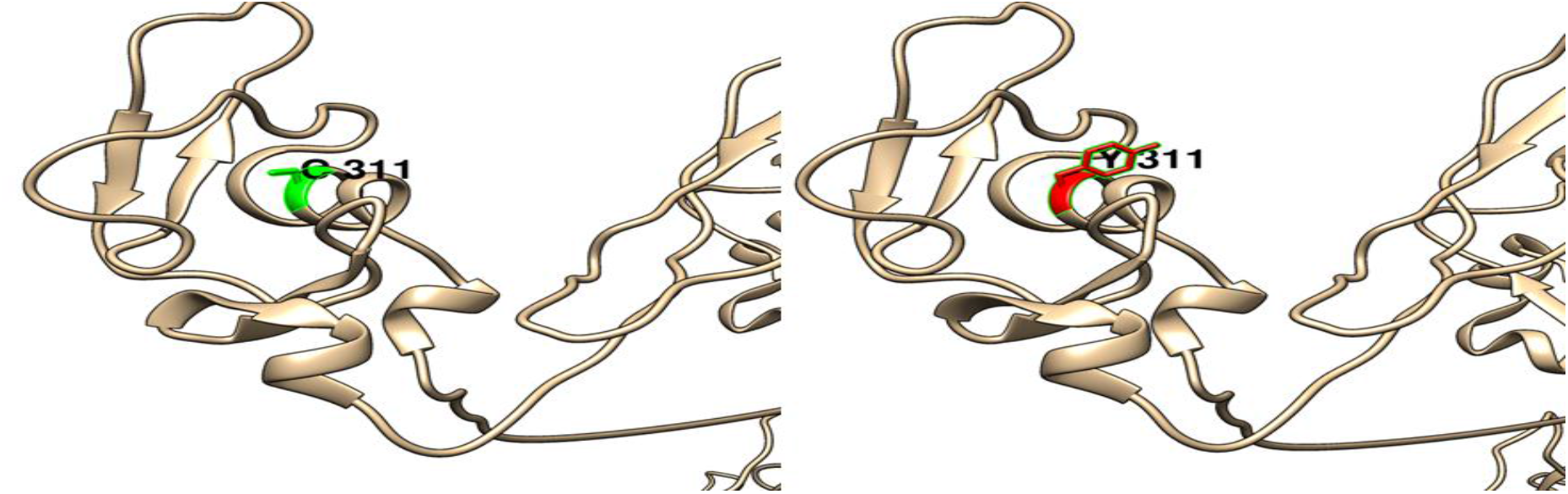
(C311Y): change in the amino Cysteine (green box) into a Tyrosine (red box) at position 311.

**Figure (10):**
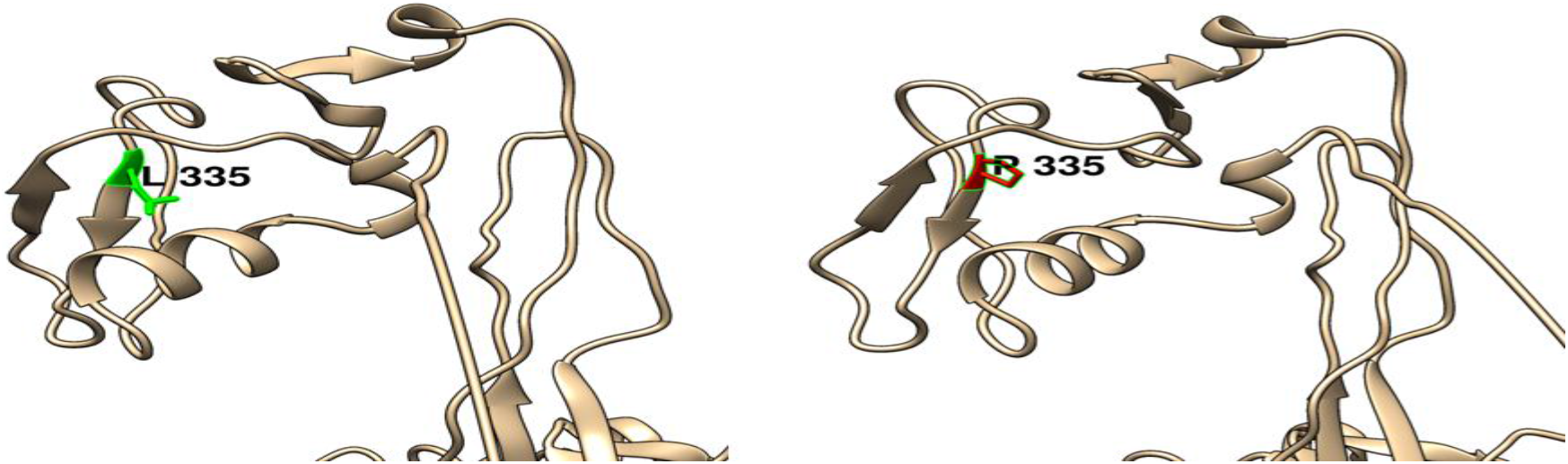
(L335P): change in the amino acid Leucine (green box) into Proline (red box) at position 335.

**Figure (11):**
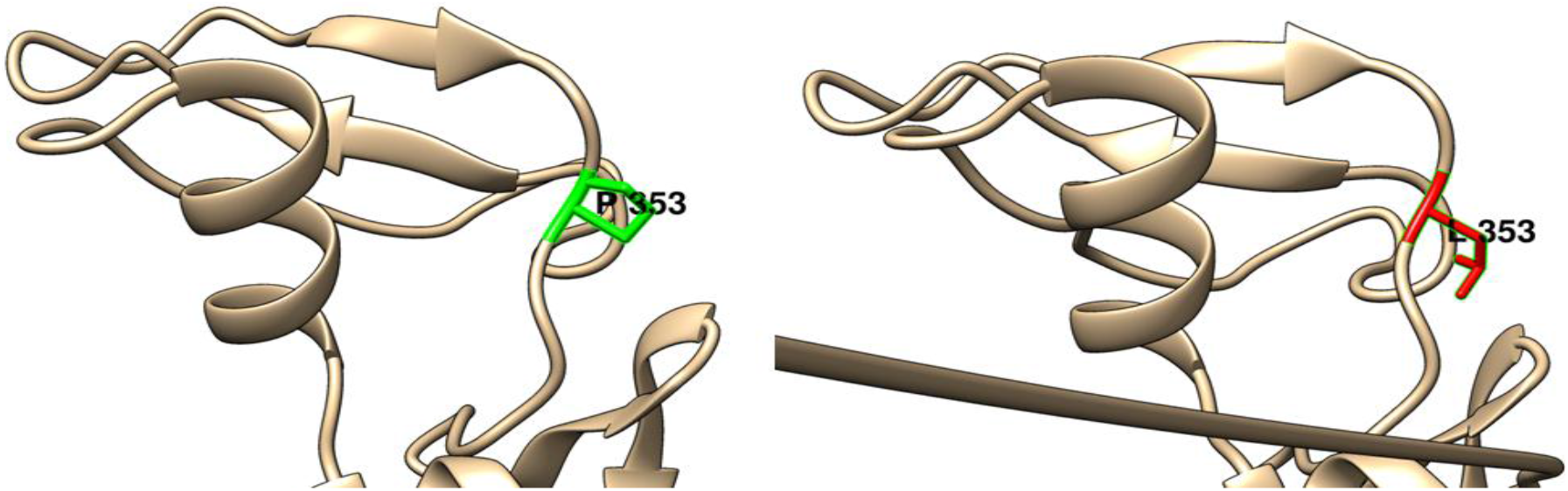
(P353L): change in the amino acid Proline (green box) into Leucine (red box) at position 353.

**Figure (12):**
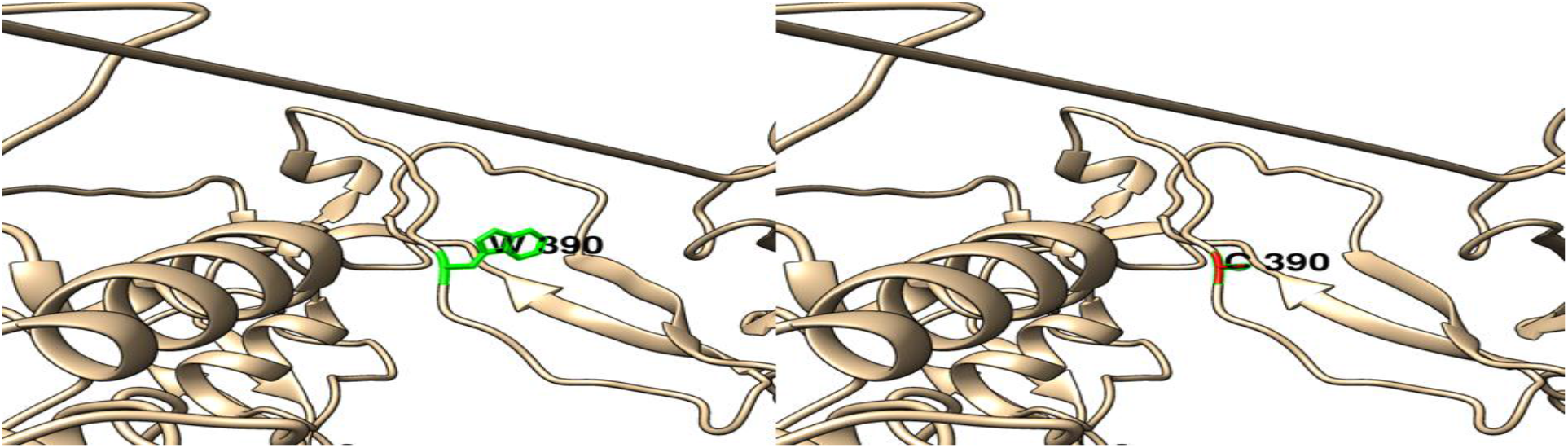
(W390C): change in the amino acid Tryptophan (green box) into Cysteine (red box) at position 390

**Figure (13):**
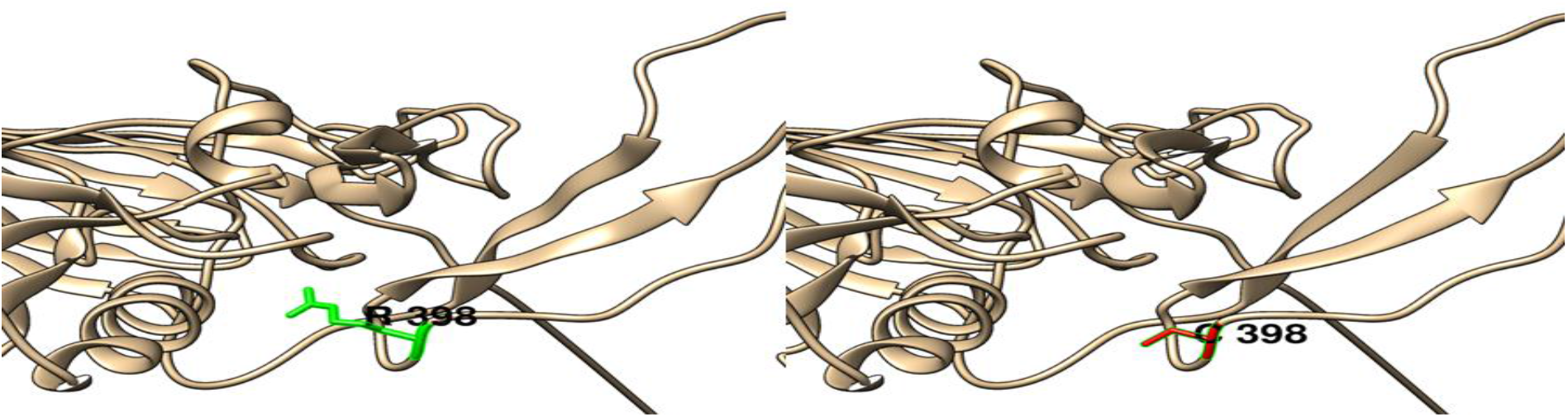
(R398C): change in the amino acid Arginine (green box) into Cysteine (red box) at position 398.

**Figure (14):**
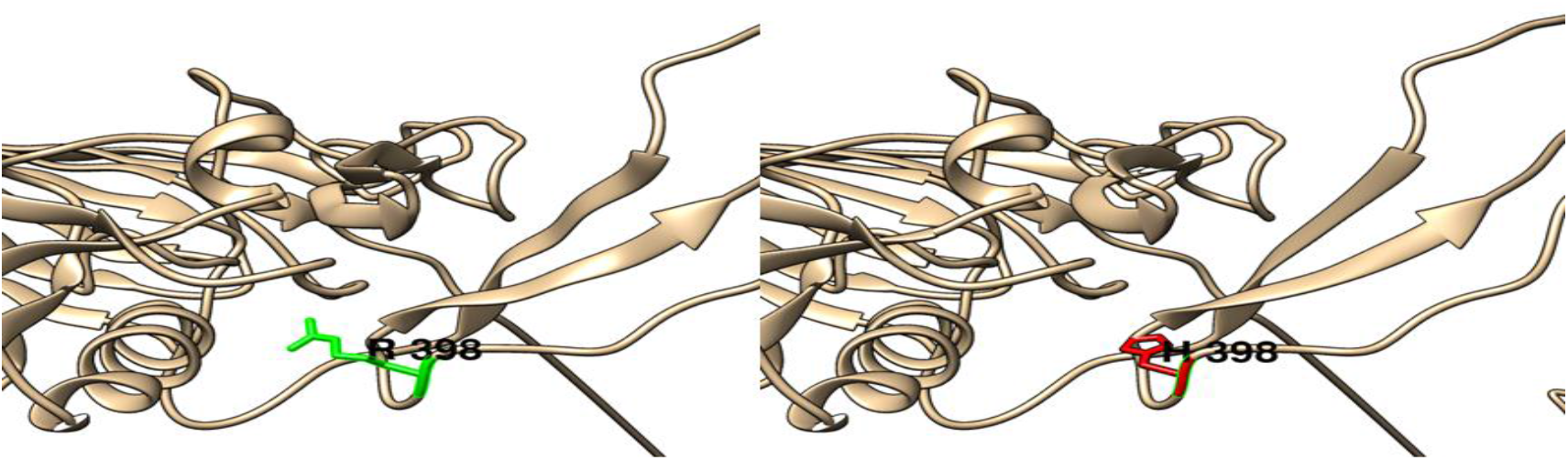
(R398H): change in the amino acid Arginine (green box) into Histidine (red box) at position 398.

**Figure (15):**
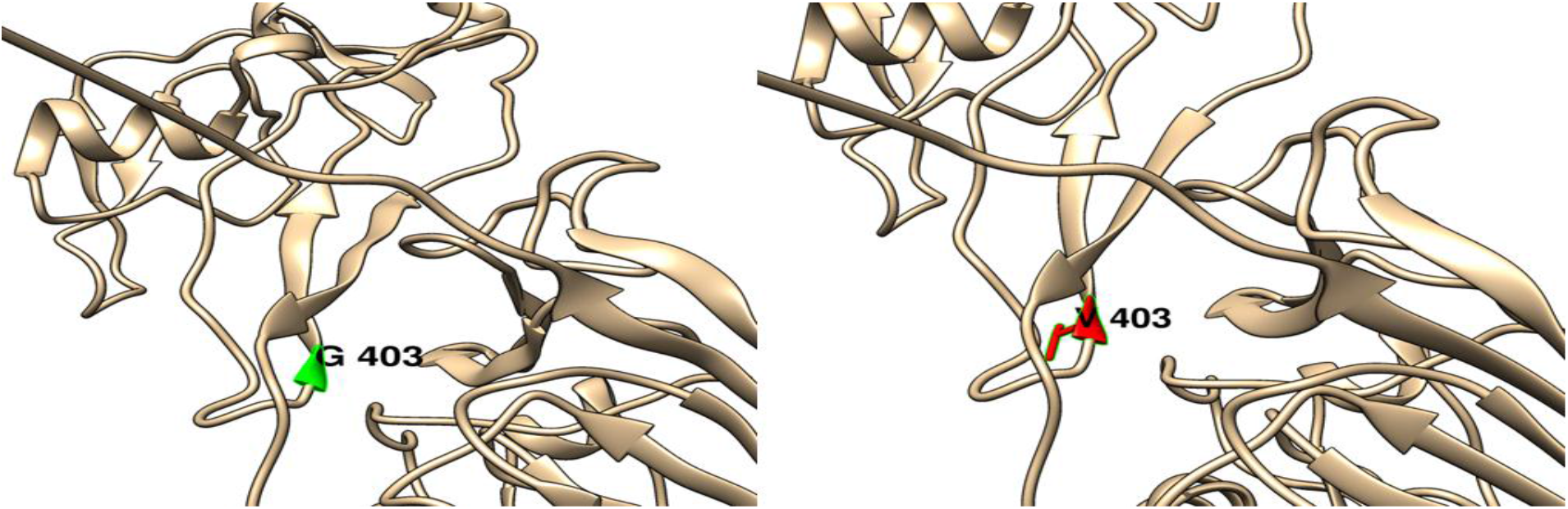
(G403V): change in the amino acid Glycine (green box) into Valine (red box) at position 403.

**Figure (16):**
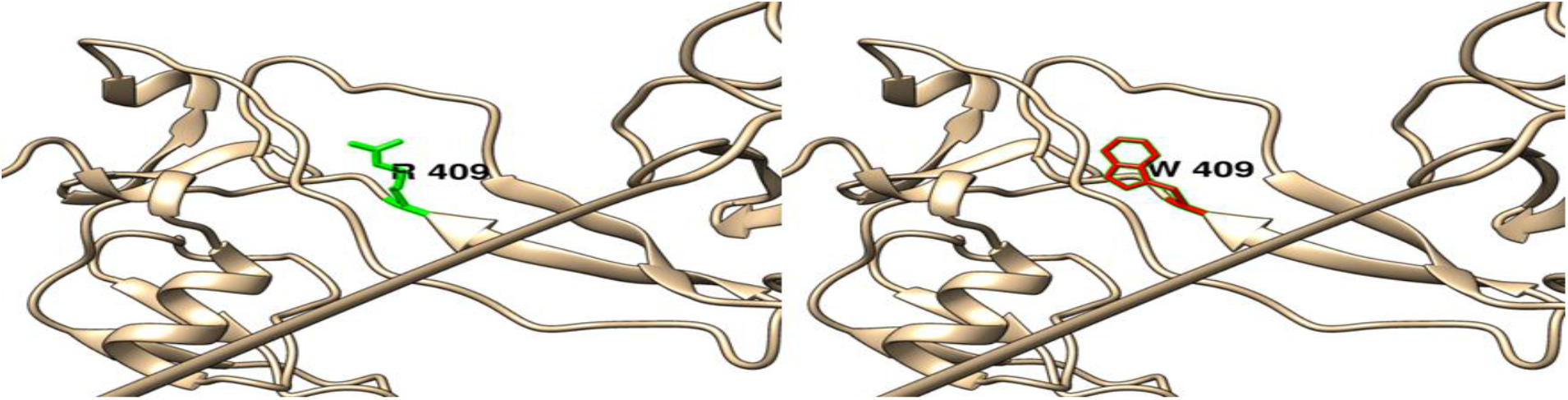
(R409W): change in the amino acid Arginine (green box) into Tryptophan (red box) at position 409.

**Figure (17):**
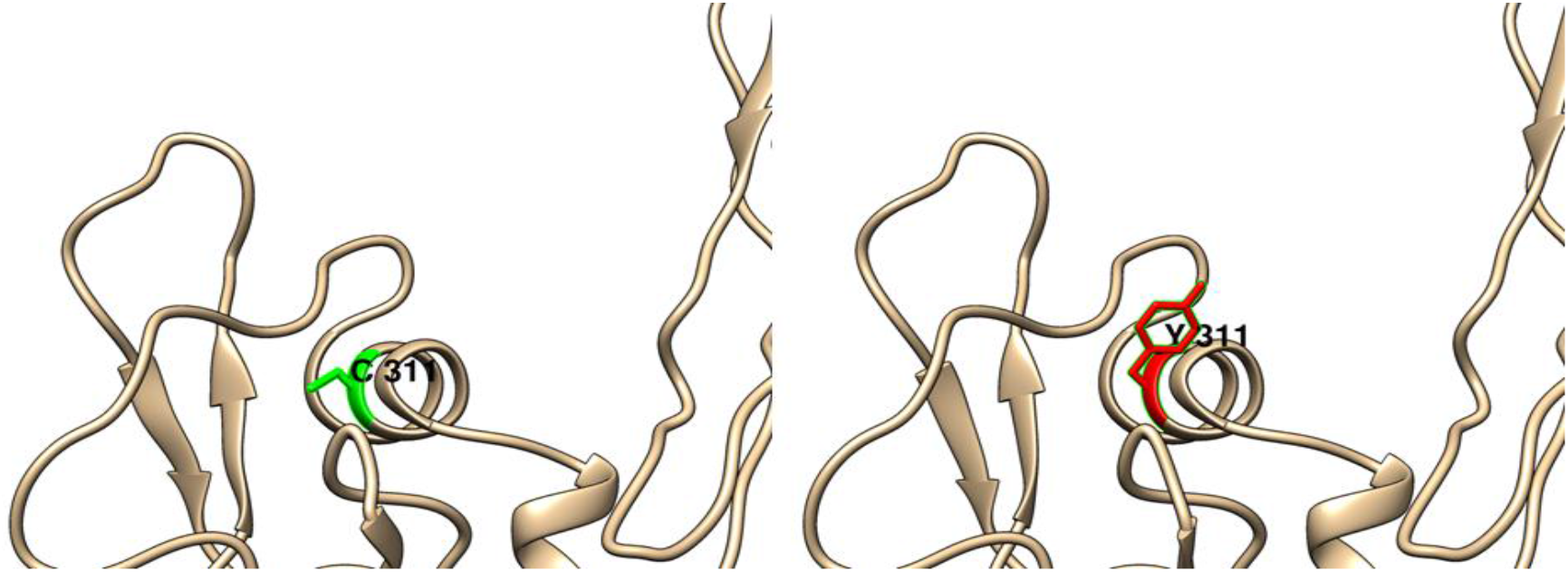
(C411Y): change in the amino acid Cysteine (green box) into Tyrosine (red box) at position 411.

**Figure (18):**
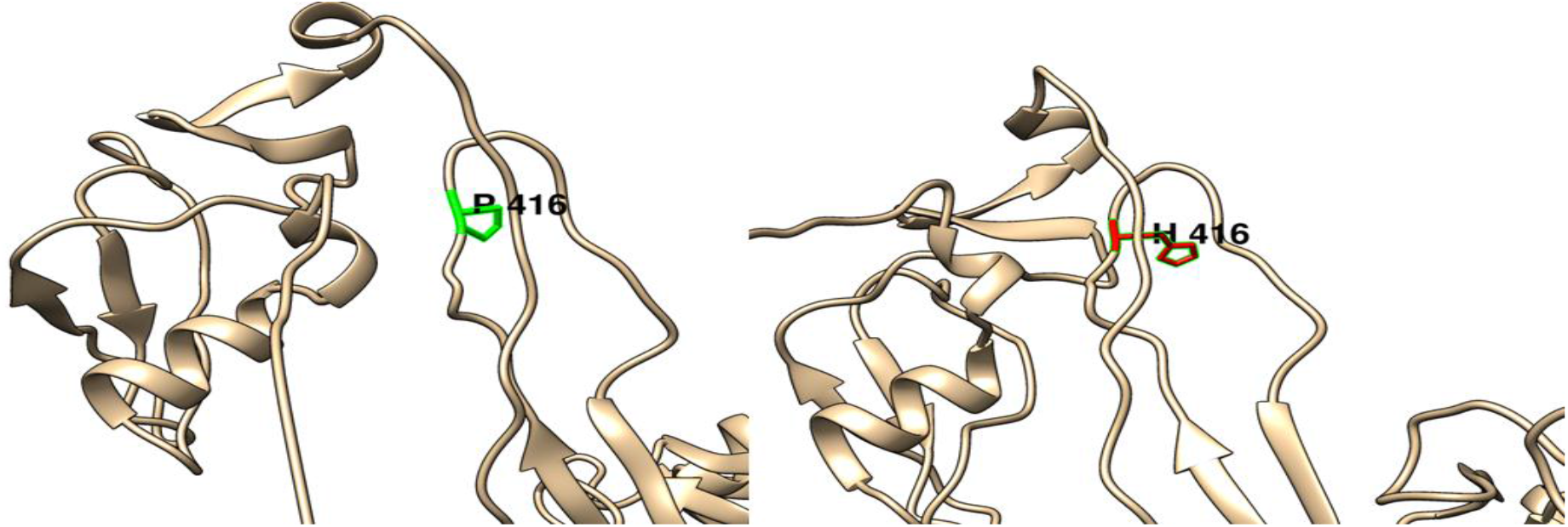
(P416H): change in the amino acid Proline (green box) into Histidine (red box) at position 416.

**Figure (19):**
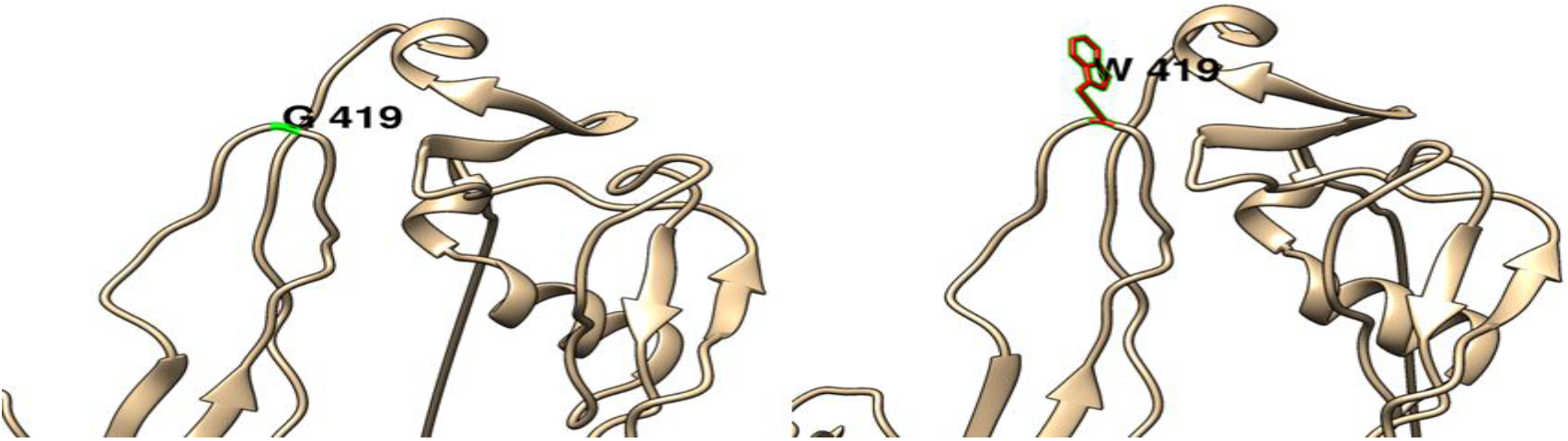
(G419W): change in the amino acid Glycine (green box) into Tryptophan (red box) at position 419.

**Figure (20):**
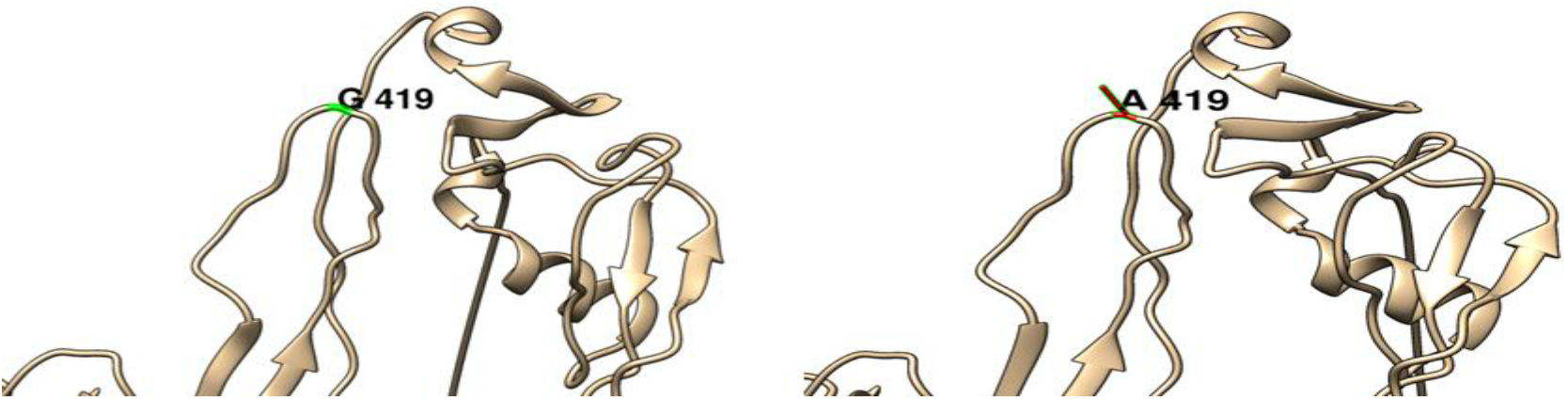
(G419A): change in the amino acid Glycine (green box) into Alanine (red box) at position 419.

**Figure (21):**
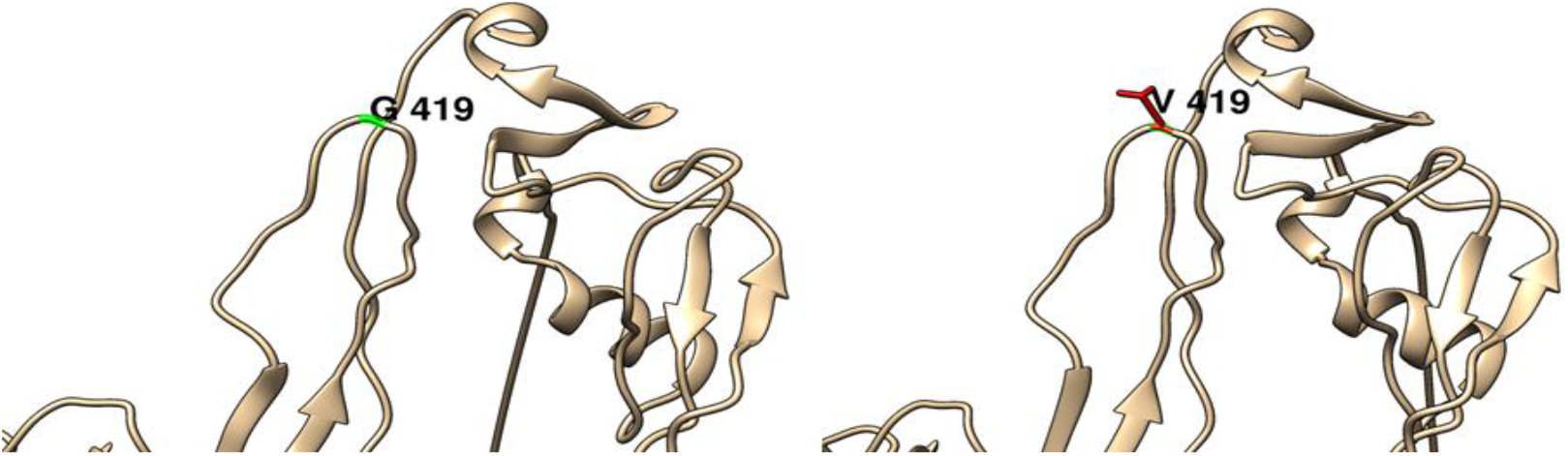
(G419V): change in the amino acid Glycine (green box) intoValine (red box) at position 419.

**Figure (22):**
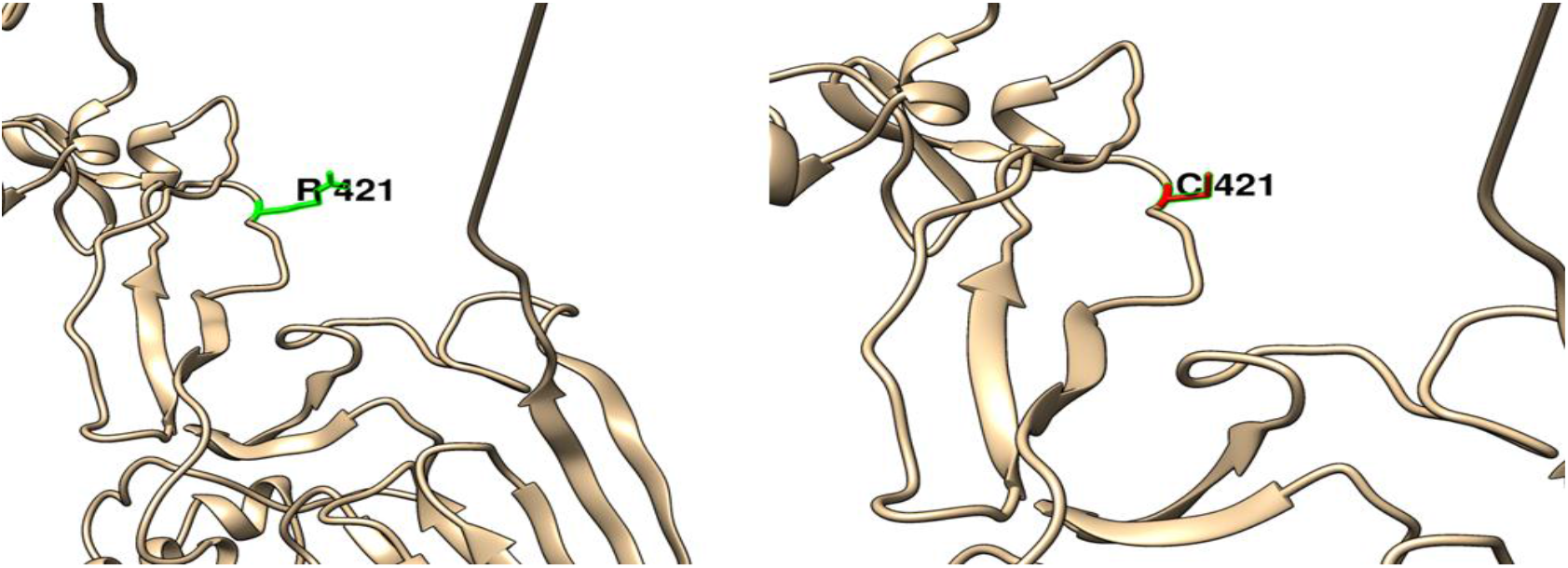
(R421C): change in the amino acid Arginine (green box) into Cysteine (red box) at position 421.

**Figure (23):**
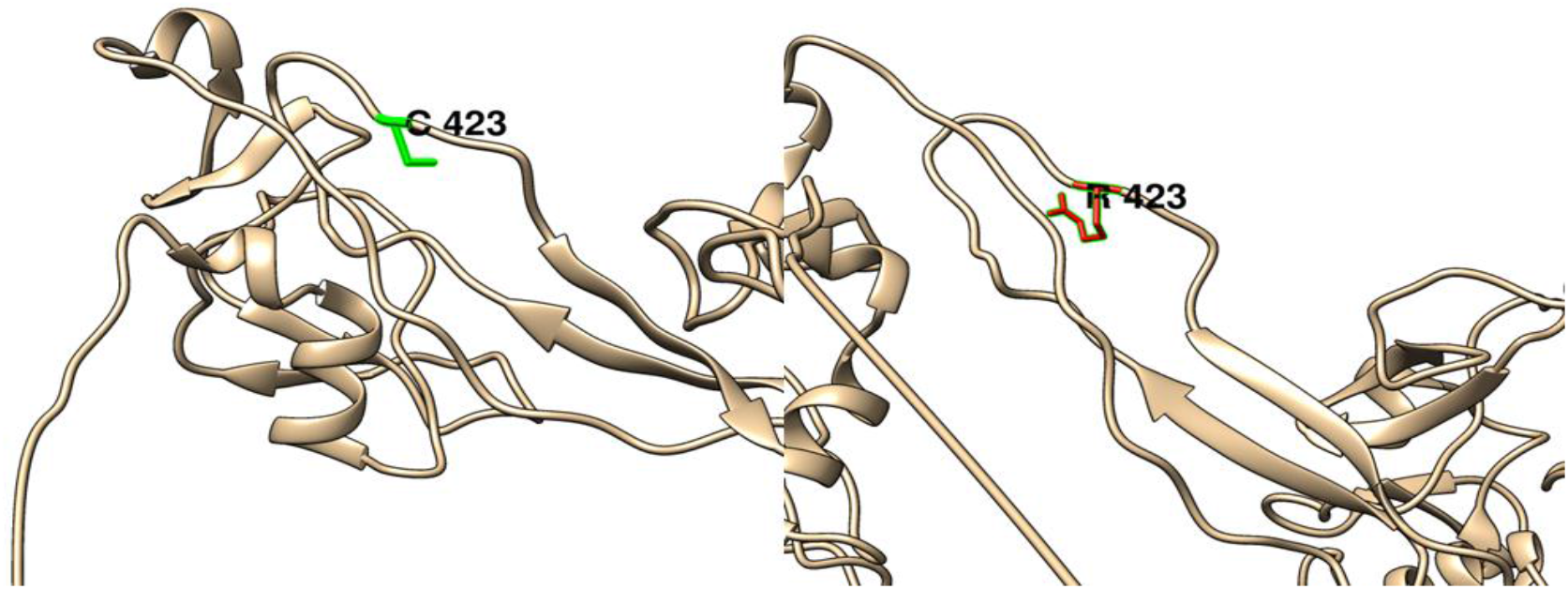
(C423R): change in the amino acid Cysteine (green box) into Arginine (red box) at position 423.

**Figure (24):**
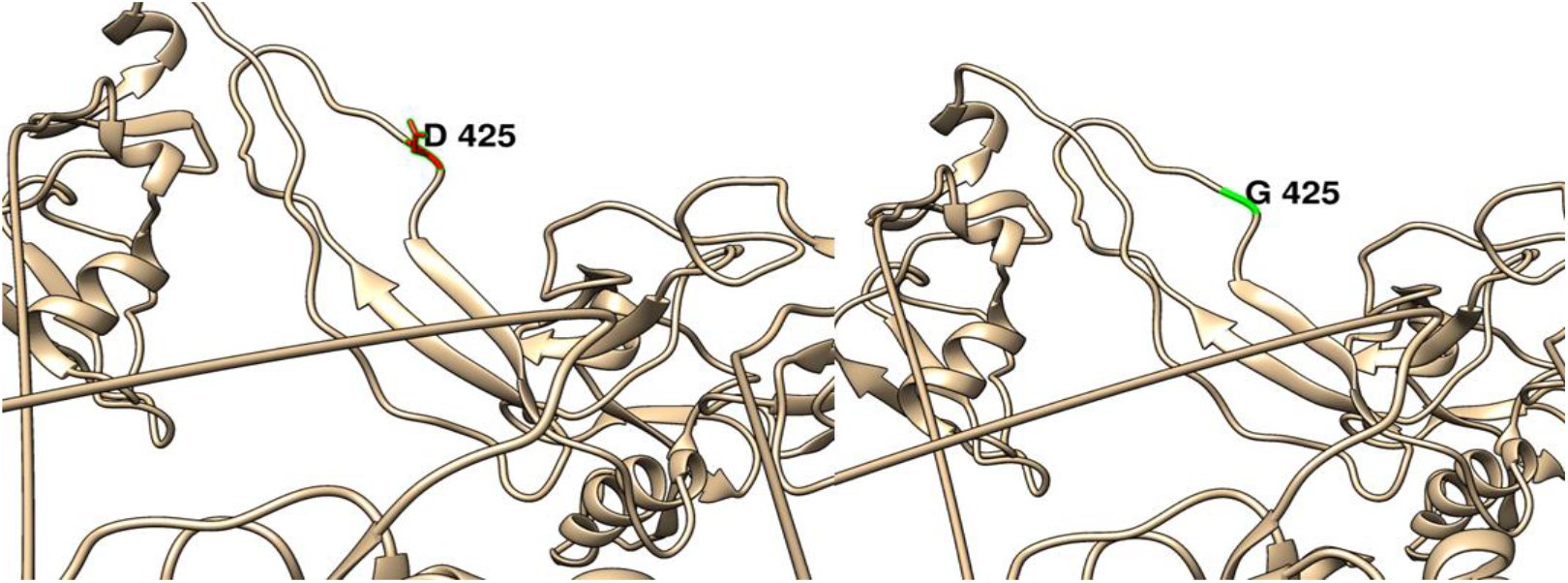
(G425D): change in the amino acid Glycine (green box) into Aspartic Acid (red box) at position 425.

**Figure (25):**
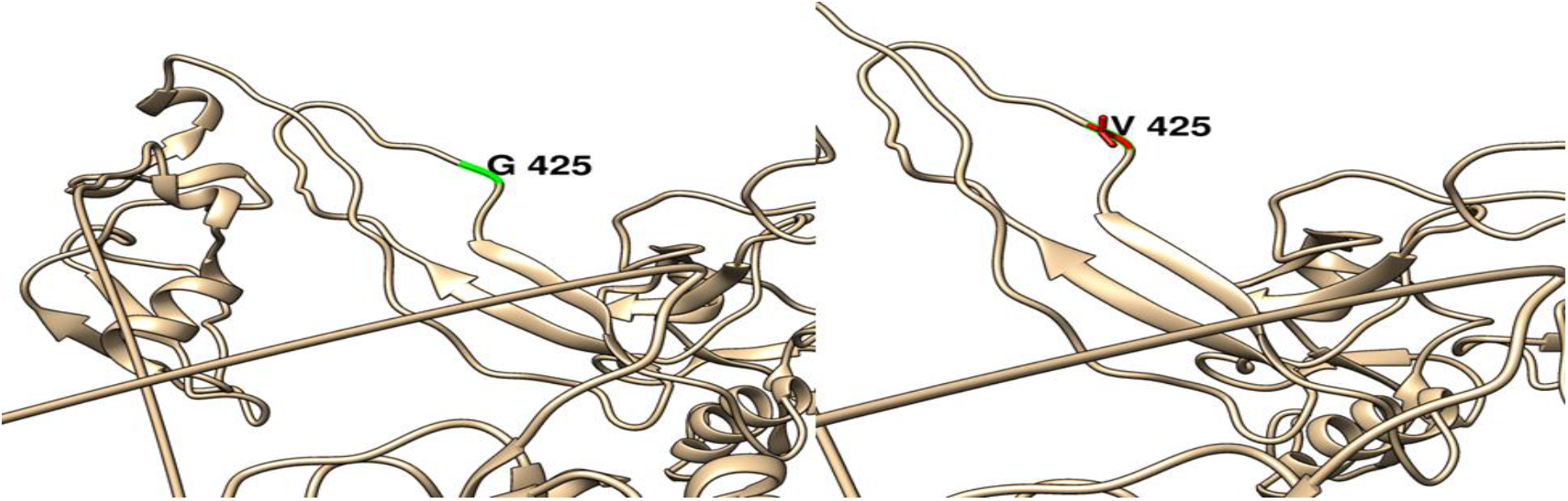
(G425V): change in the amino acid Glycine (green box) into Valine (red box) at position 425.

**Figure (26):**
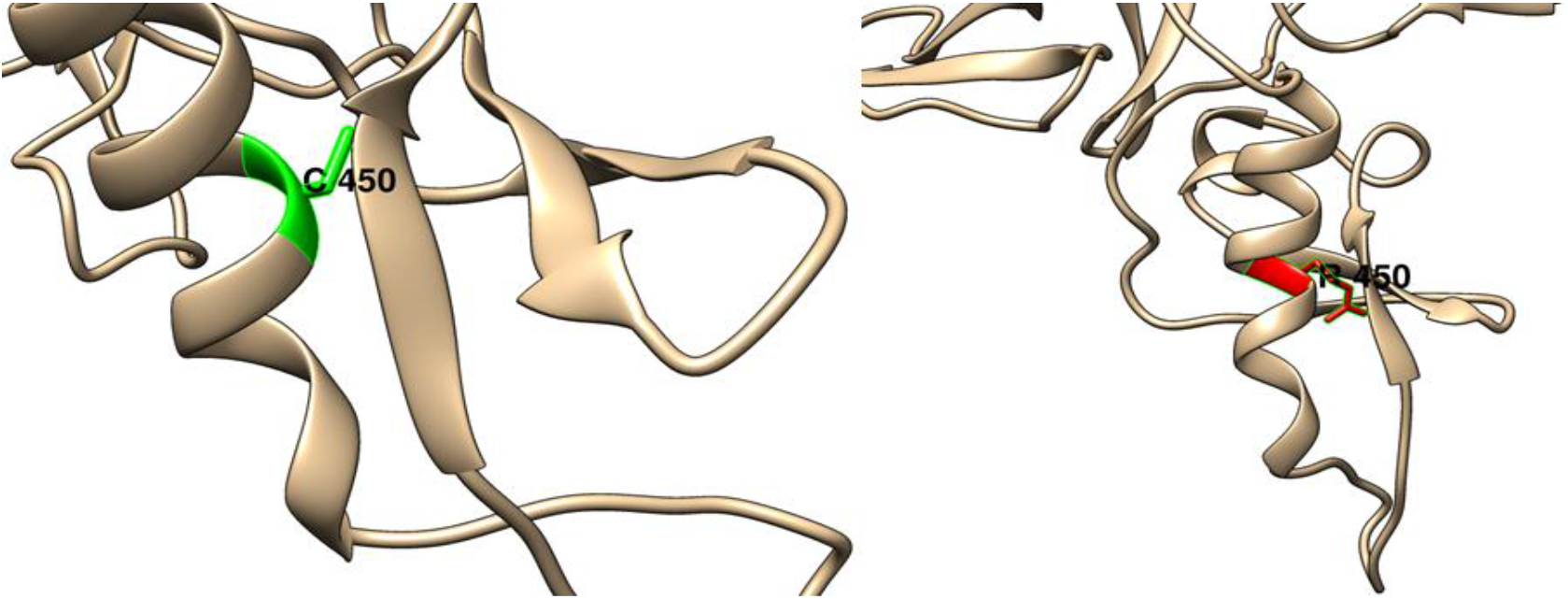
(C450R): change in the amino acid Cysteine (green box) into Arginine (red box) at position 450.

**Figure (27):**
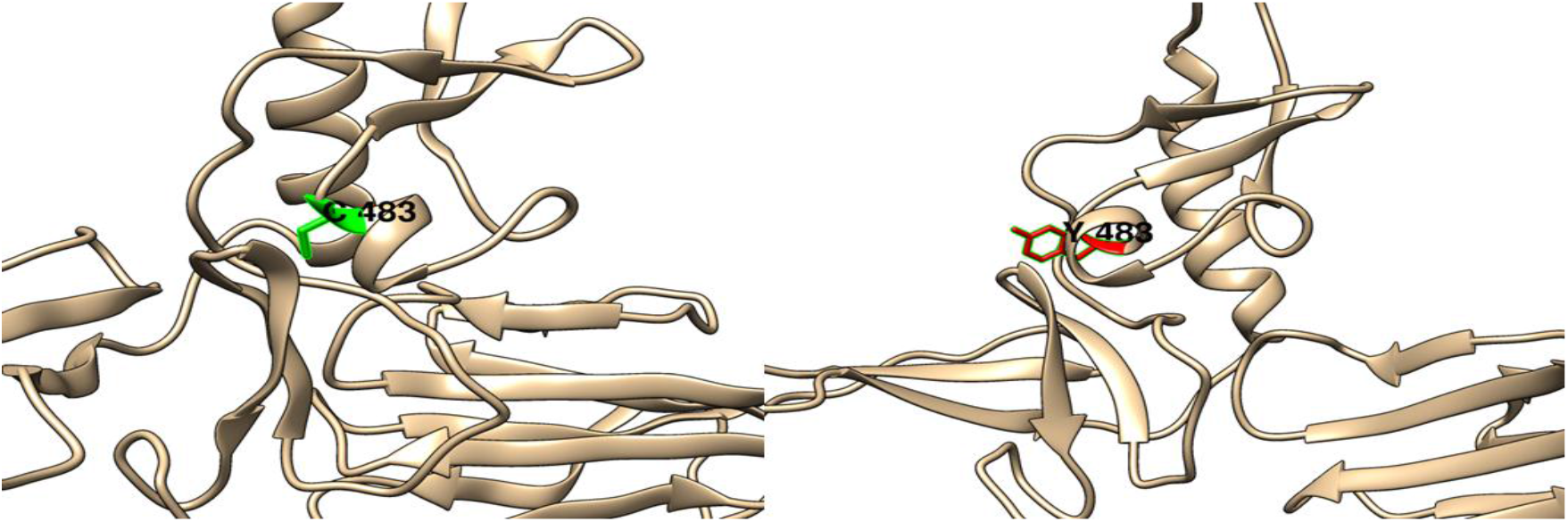
(C483Y): change in the amino acid Cysteine (green box) into Tyrosine (red box) at position 483.

**Figure (28):**
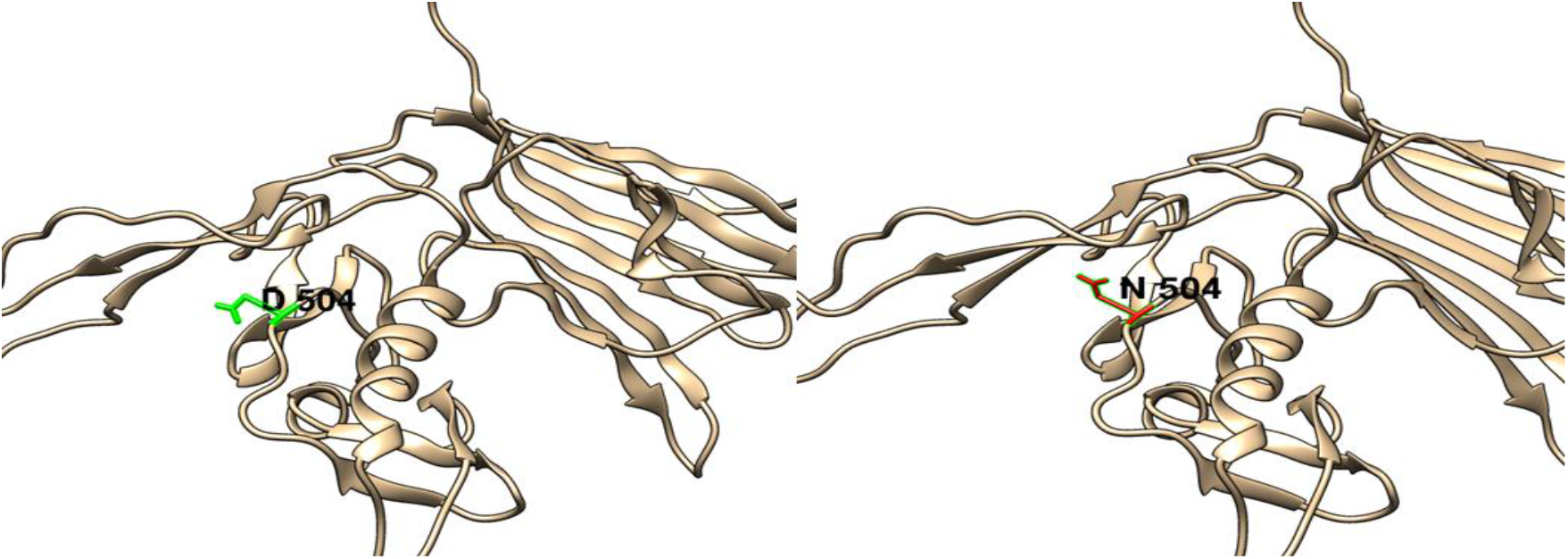
(D504N): change in the amino acid Aspartic Acid (green box) into Asparagine (red box) at position 504.

**Figure (29):**
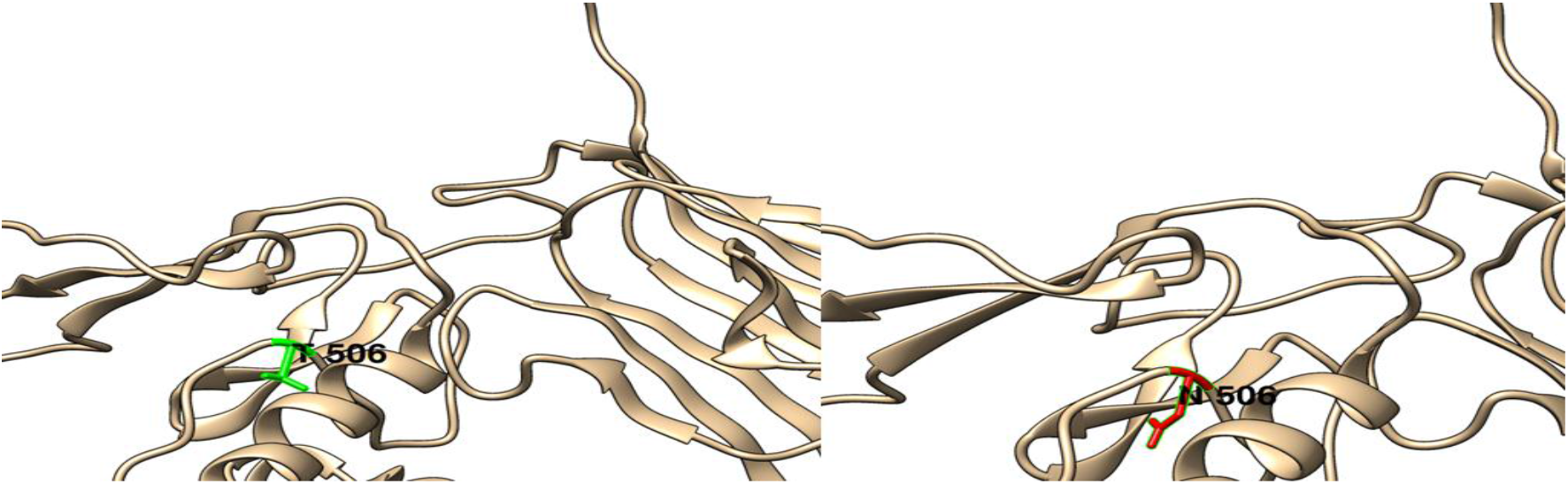
(T506N): change in the amino acid Threonine (green box) into Asparagine (red box) at position 506.

**Figure (30):**
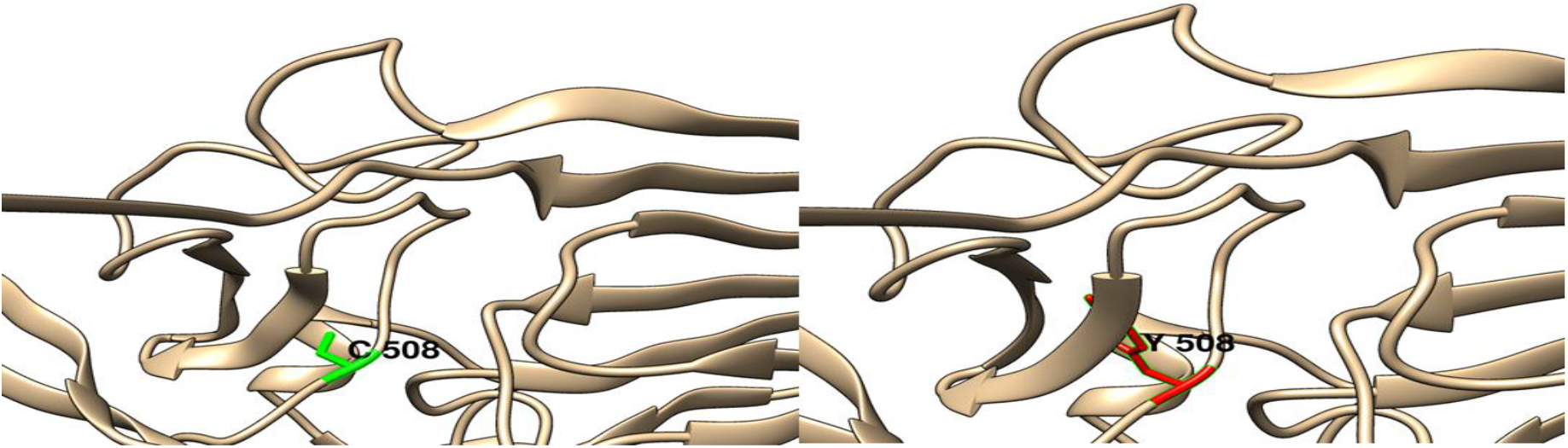
(C508Y): change in the amino acid Cysteine (green box) into Tyrosine (red box) at position 508.

**Figure (31):**
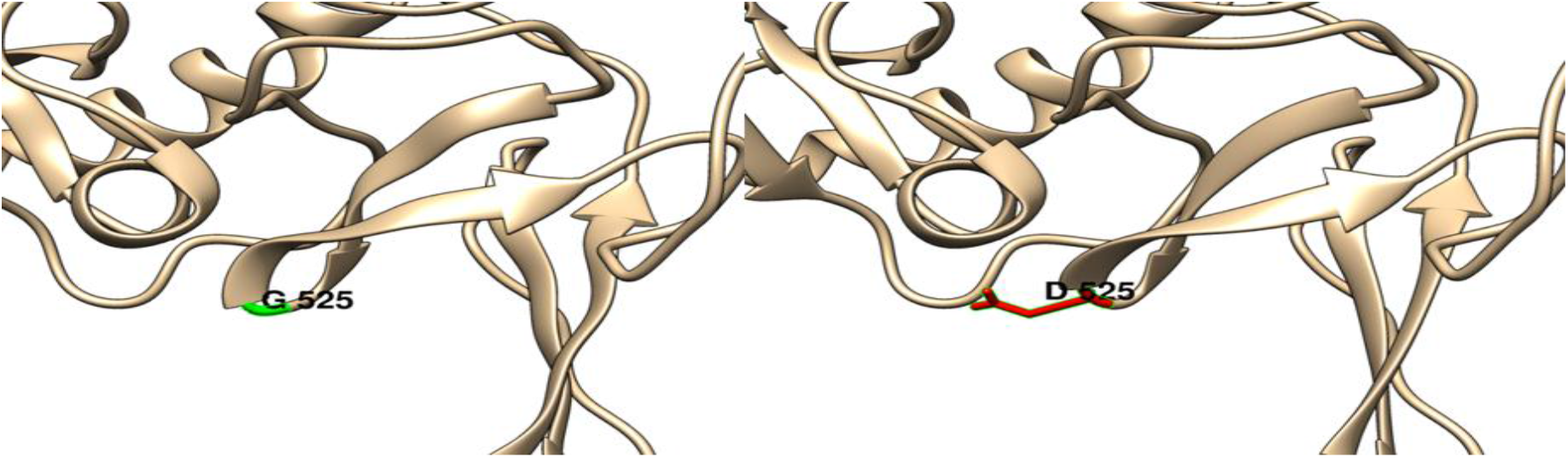
(G525D): change in the amino acid Glycine (green box) into Aspartic Acid (red box) at position 525.

**Figure (32):**
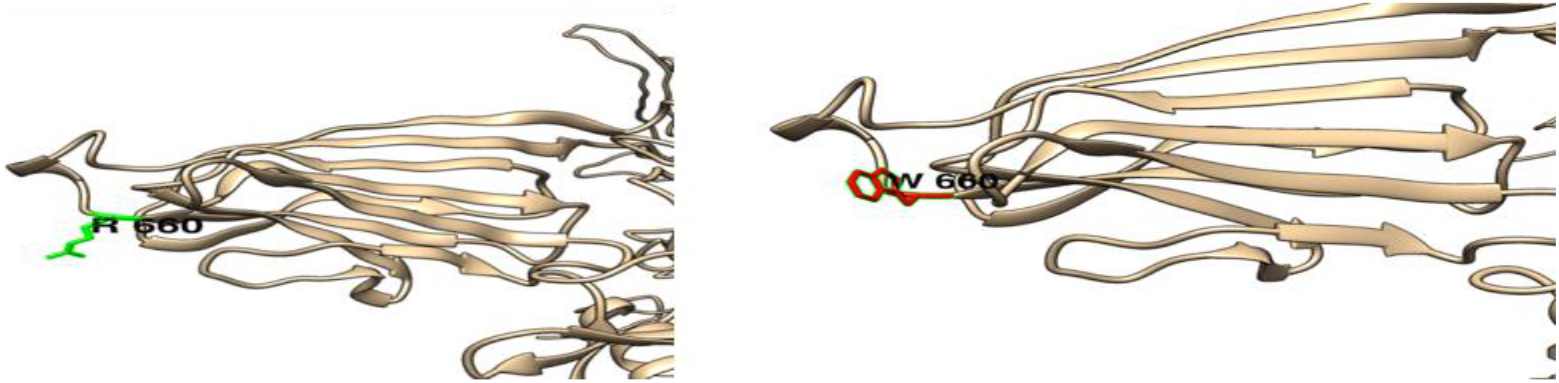
(R660W): change in the amino acid Arginine (green box) into Tryptophan (red box) at position 660.

**Figure (33):**
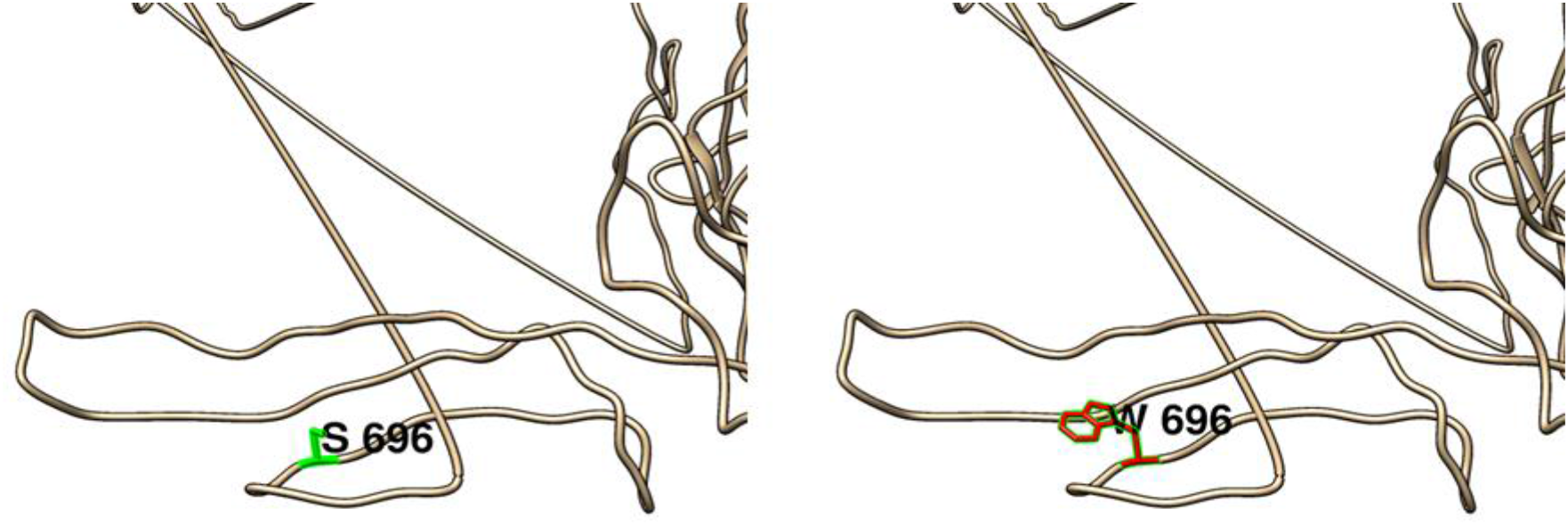
(S696W): change in the amino acid Serine (green box) into Tryptophan (red box) at position 696.

**Figure (34):**
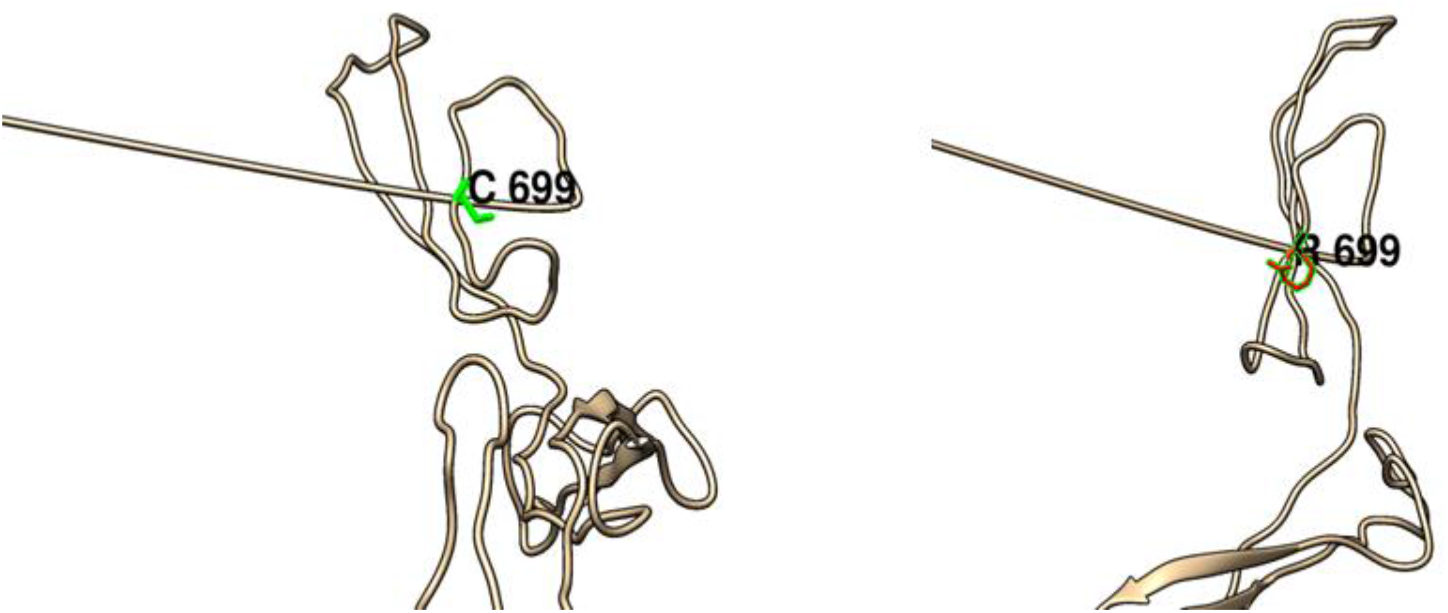
(C699R): change in the amino acid Cysteine (green box) into Arginine (red box) at position 699.

**Figure (35):**
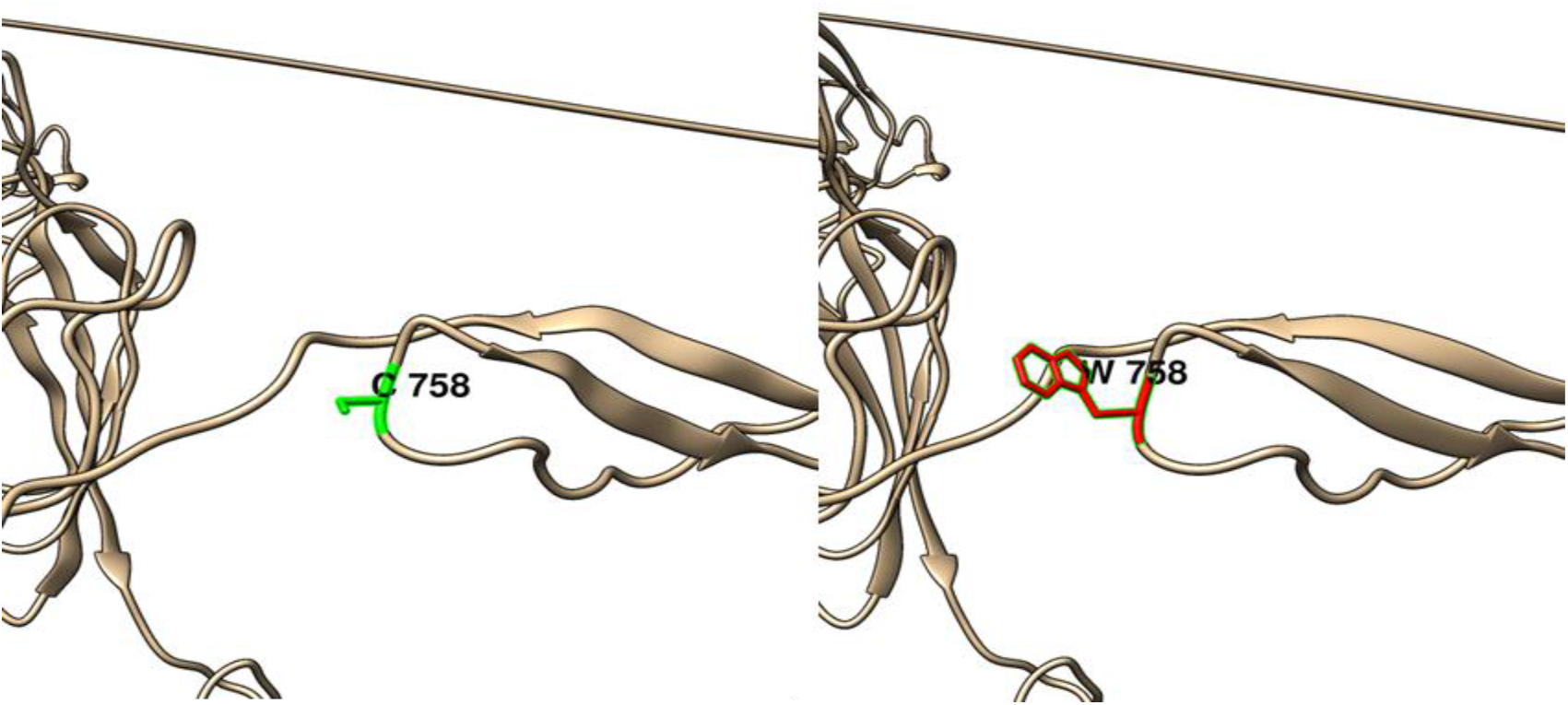
(C758W): change in the amino acid Cysteine (green box) into Tryptophan (red box) at position 758.

**Figure (36):**
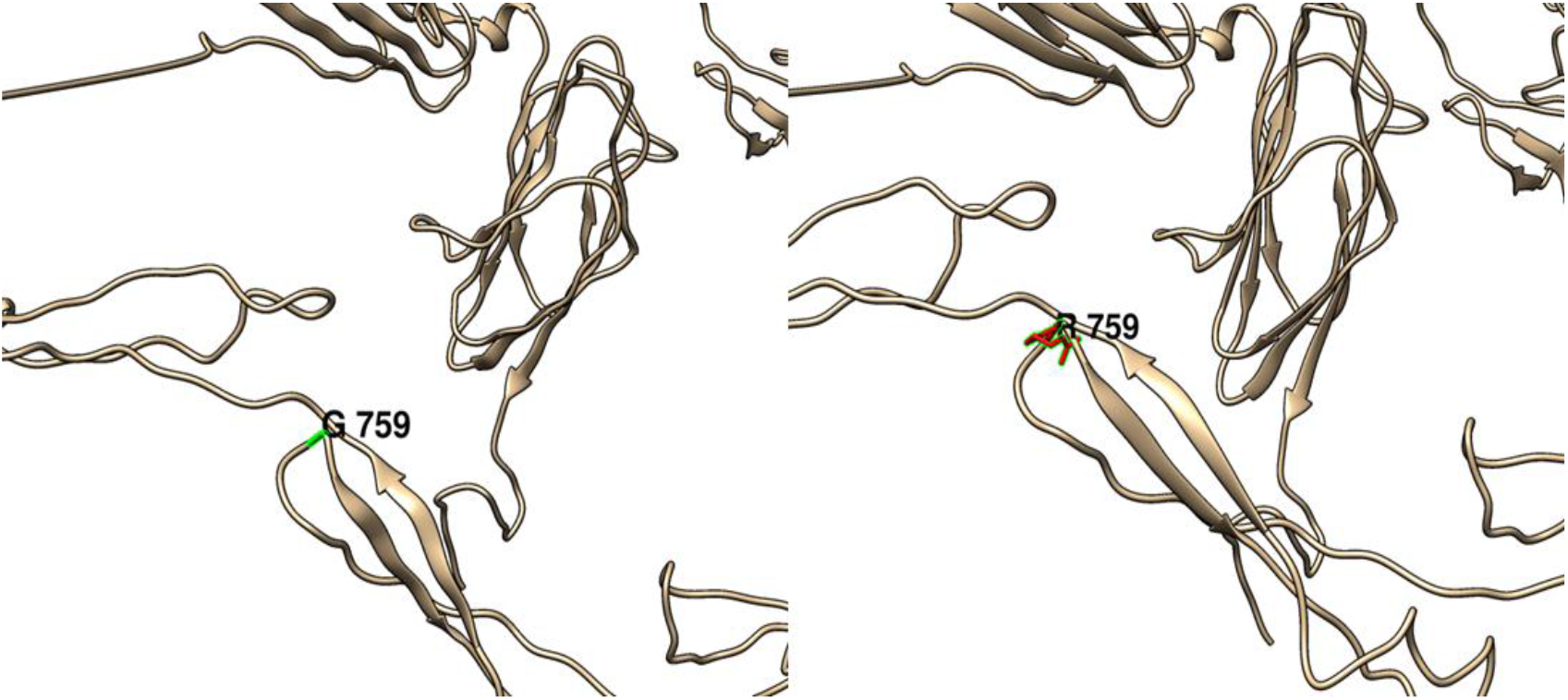
(G759R): change in the amino acid Glycine (green box) into Tryptophan (red box) at position 759.

**Figure (37):**
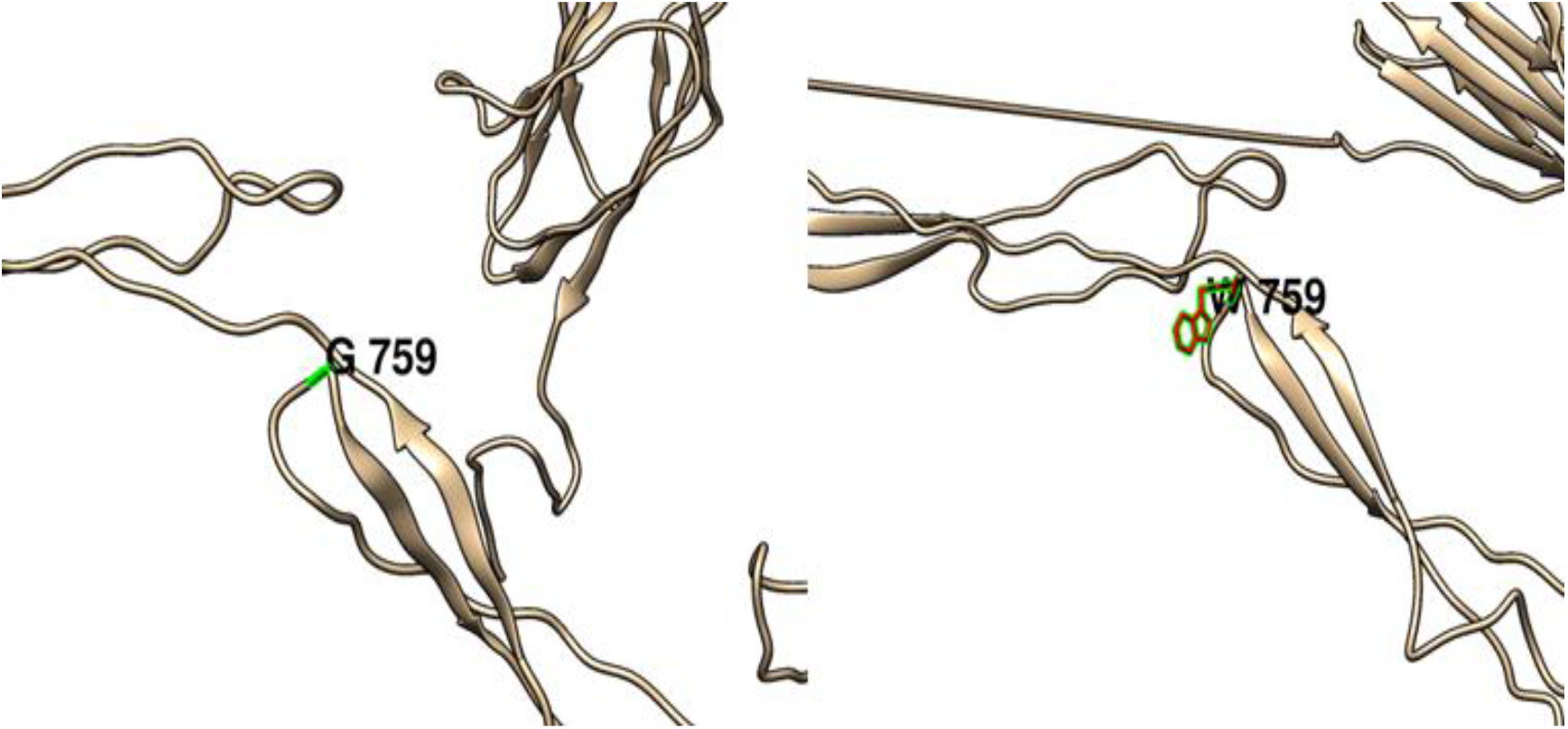
(G759W): change in the amino acid Glycine (green box) into Tryptophan (red box) at position 759.

**Figure (38):**
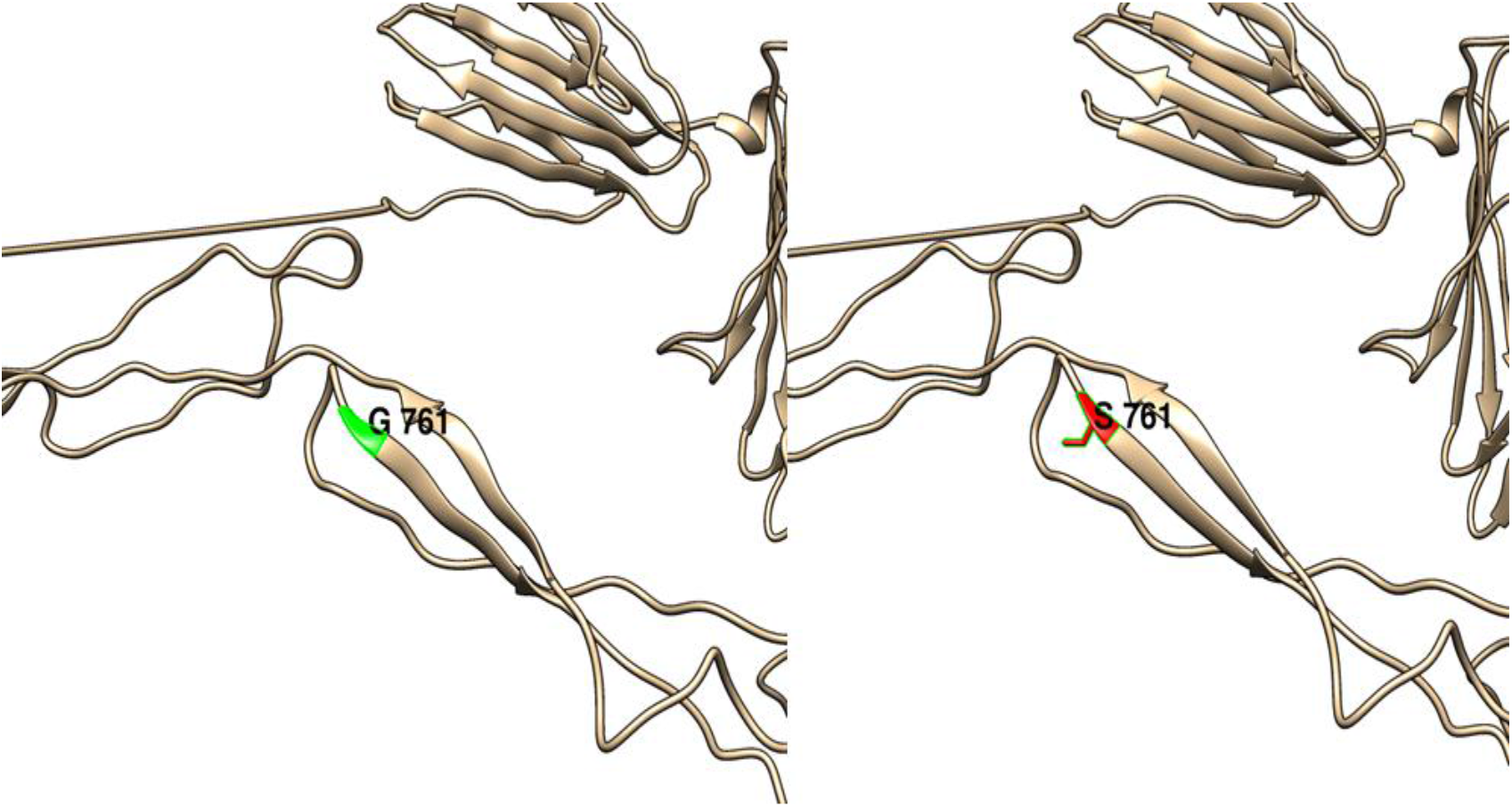
(G761S): change in the amino acid Glycine (green box) into Serine (red box) at position 761.

**Figure (39):**
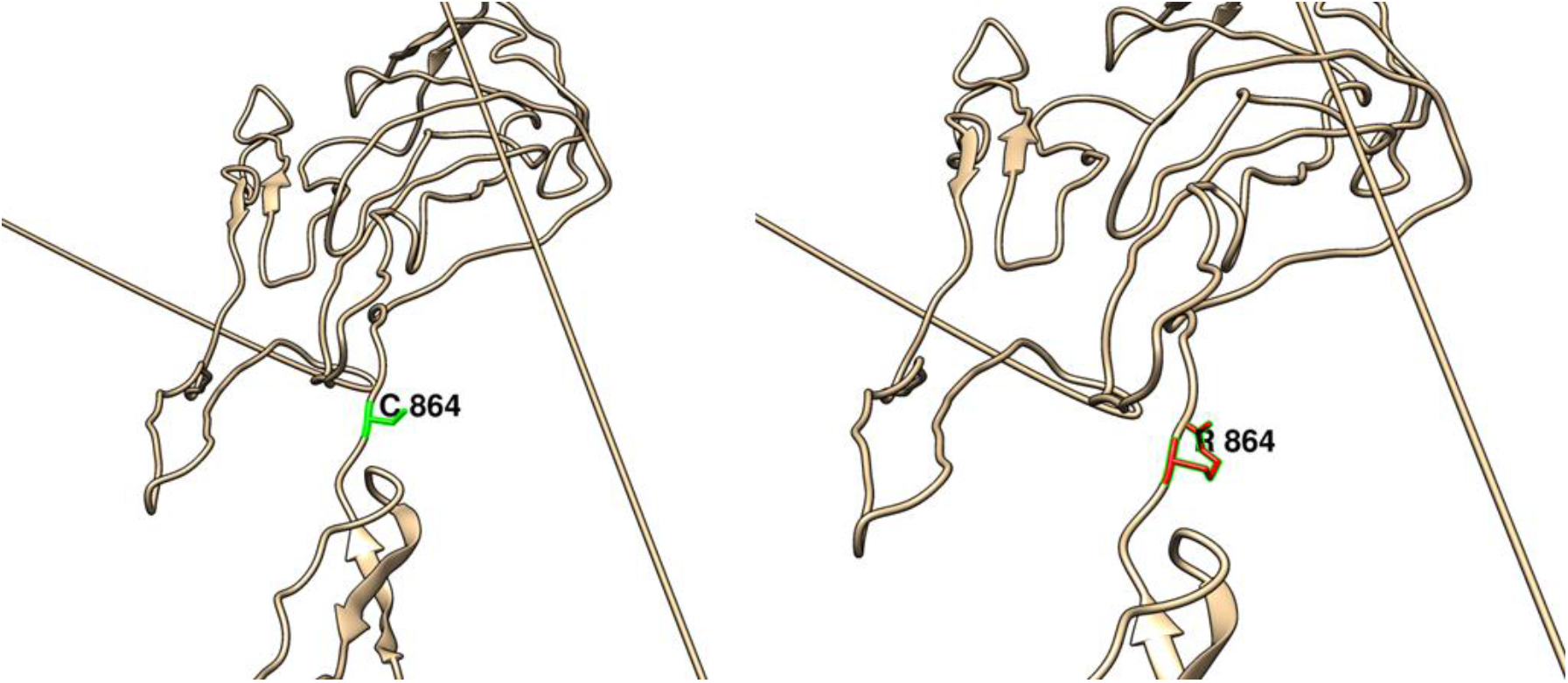
(C864R): change in the amino acid Cysteine (green box) into Arginine (red box) at position 864.

**Figure (40):**
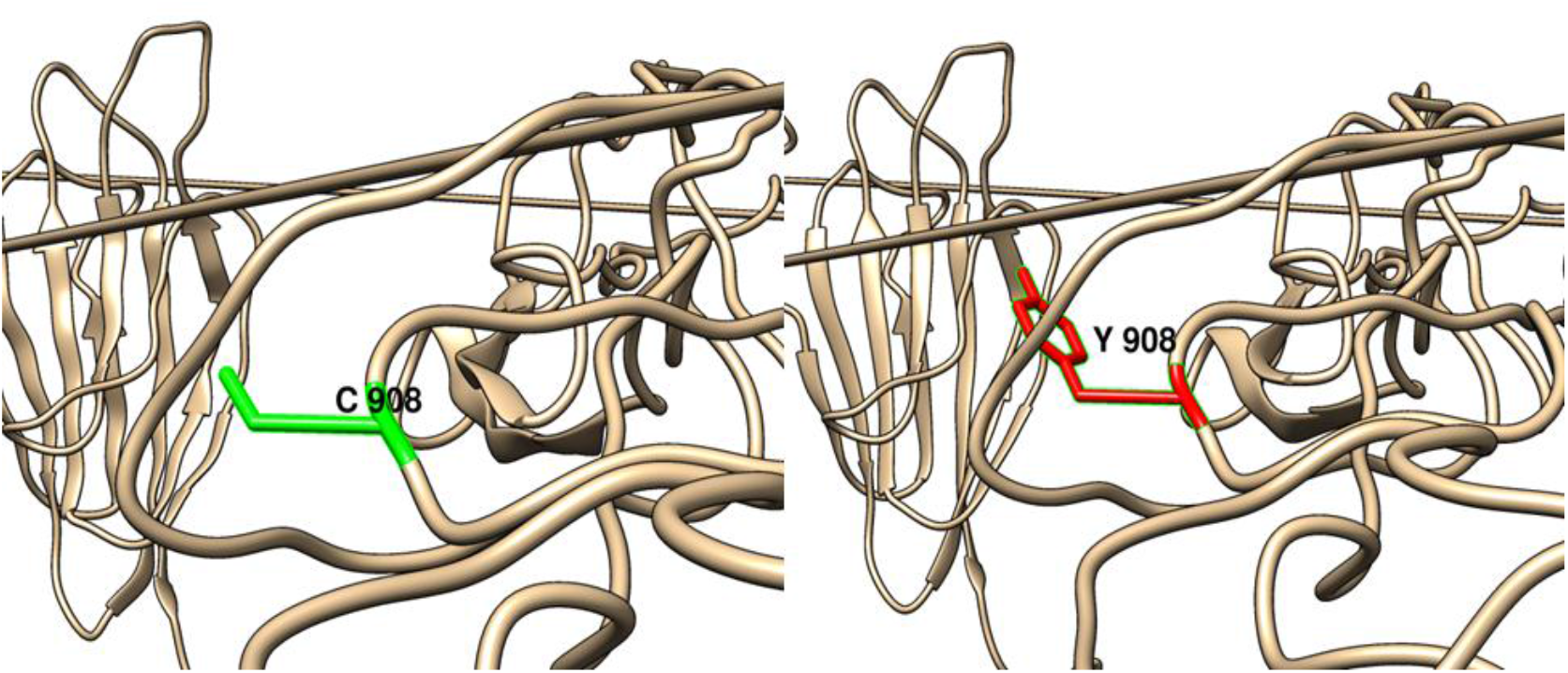
(C908Y): change in the amino acid Cysteine (green box) into Tyrosine (red box) at position 908.

**Figure (41):**
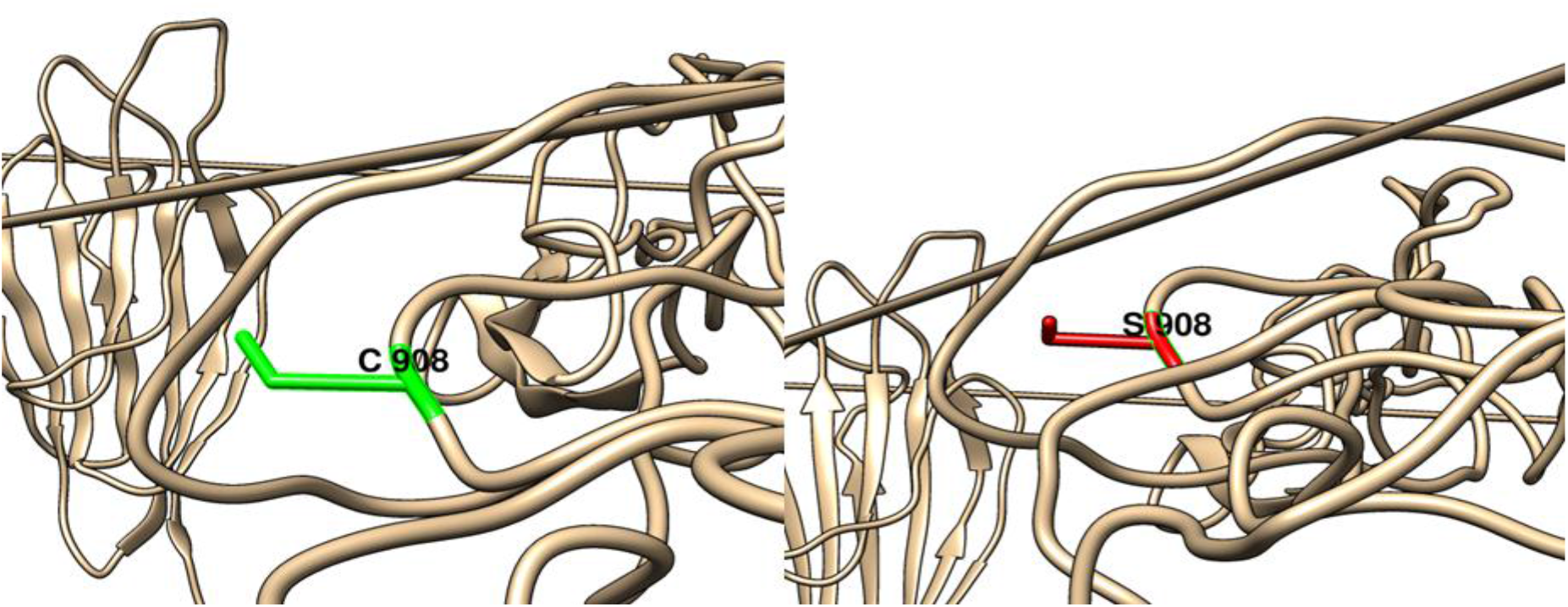
(C908S): change in the amino acid Cysteine (green box) into Serine (red box) at position 908.

**Figure (42):**
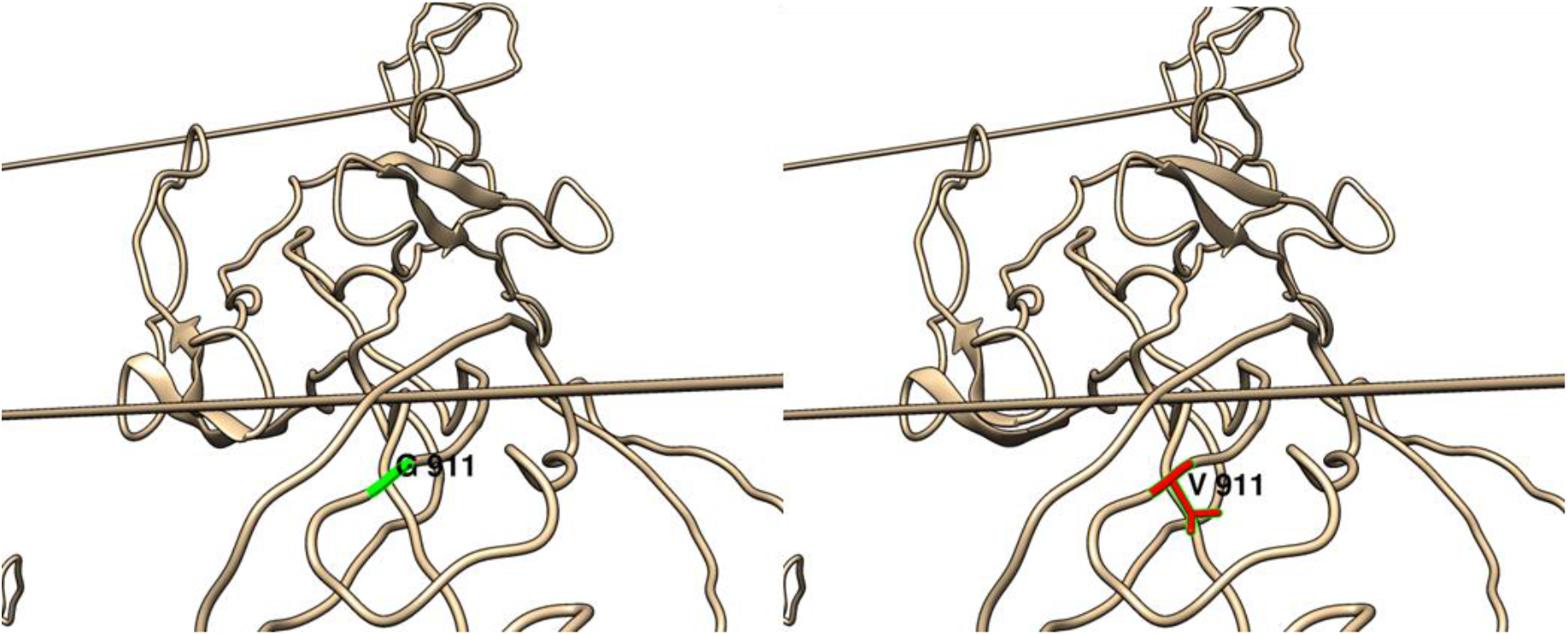
(G911V): change in the amino acid Glycine (green box) into Valine (red box) at position 911.

**Figure (43):**
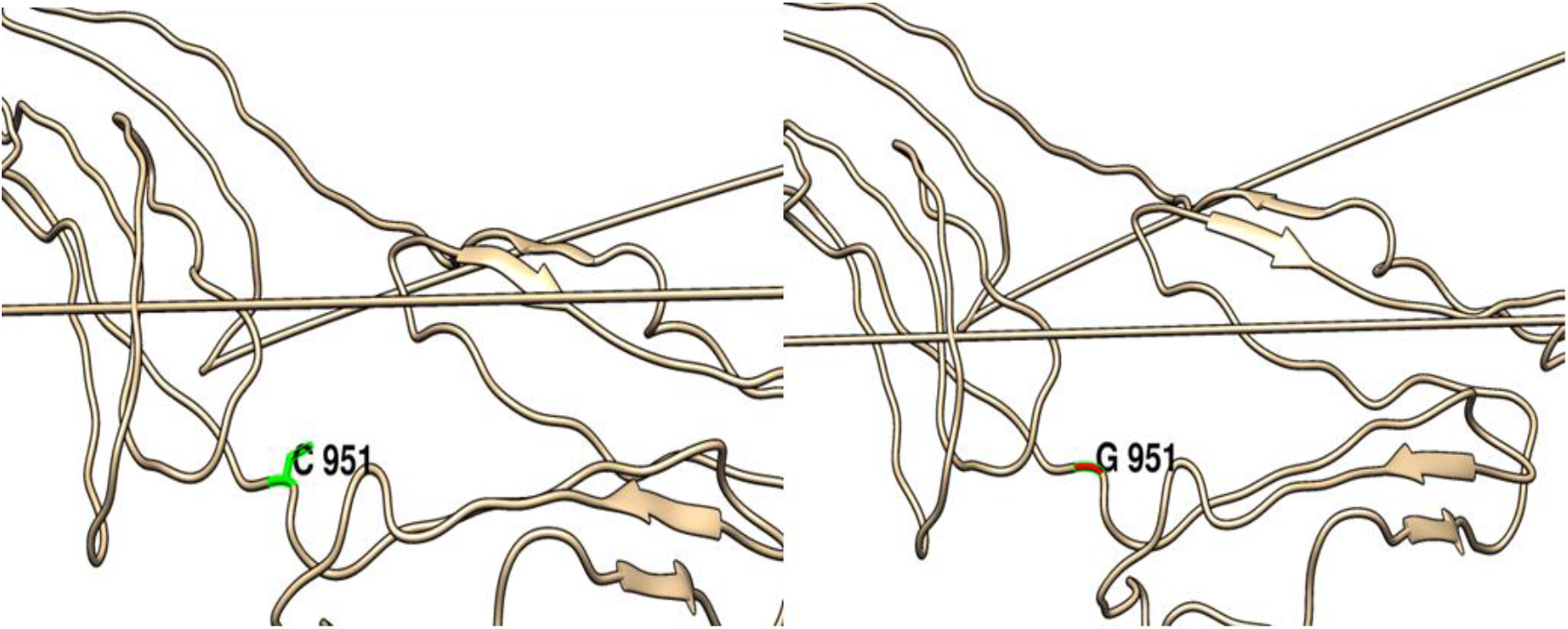
(C951G): change in the amino acid Cysteine (green box) into Glycine (red box) at position 951.

**Figure (44):**
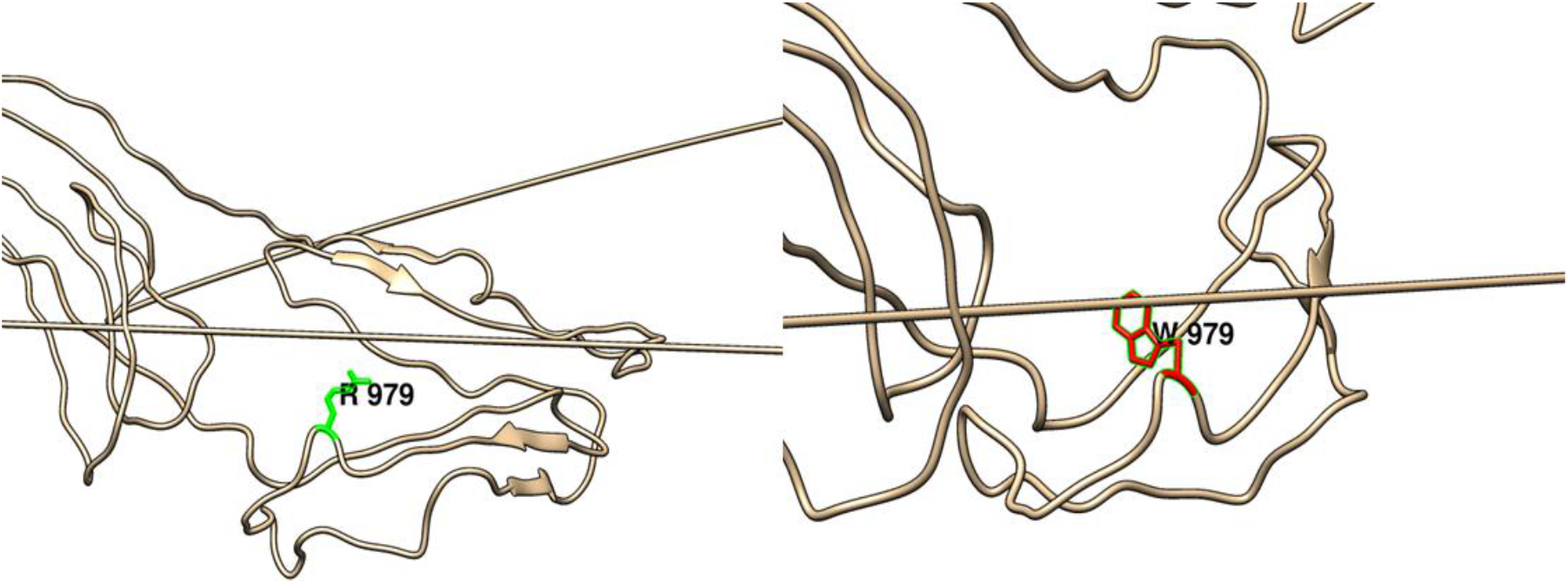
(R979W): change in the amino acid Arginine (green box) into Tryptophan (red box) at position 979.

**Figure (45):**
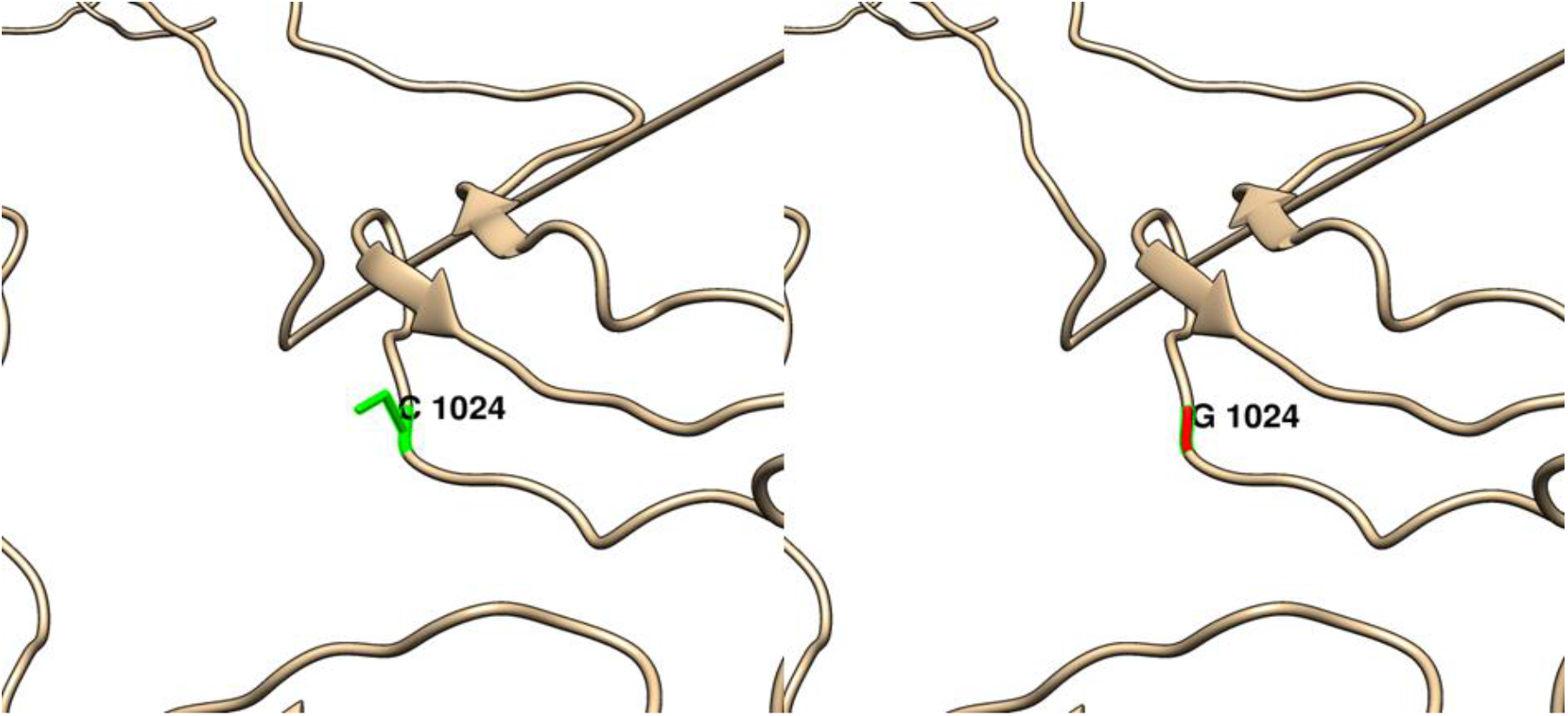
(C1024G): change in the amino acid Cysteine (green box) into Glycine (red box) at position 1024.

**Figure (46):**
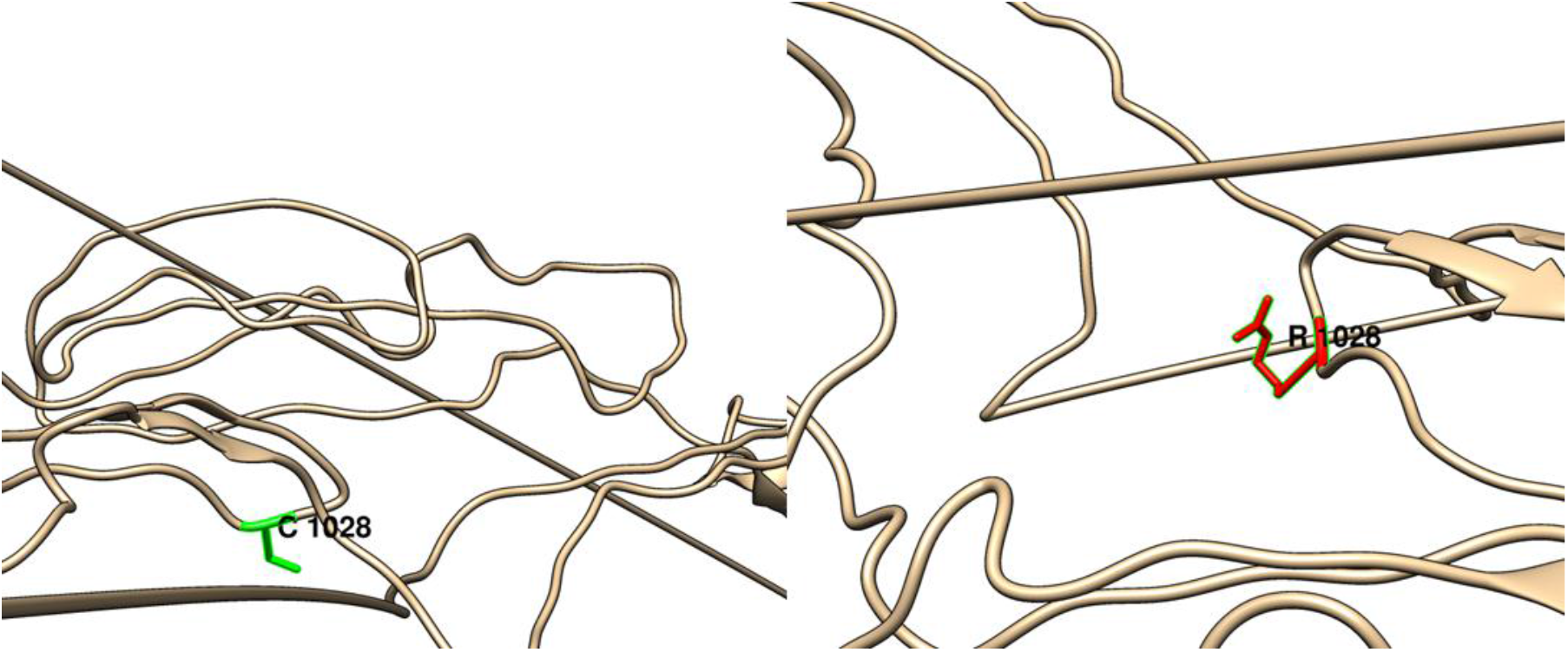
(C1028R): change in the amino acid Cysteine (green box) into Arginine (red box) at position 1028.

**Figure (47):**
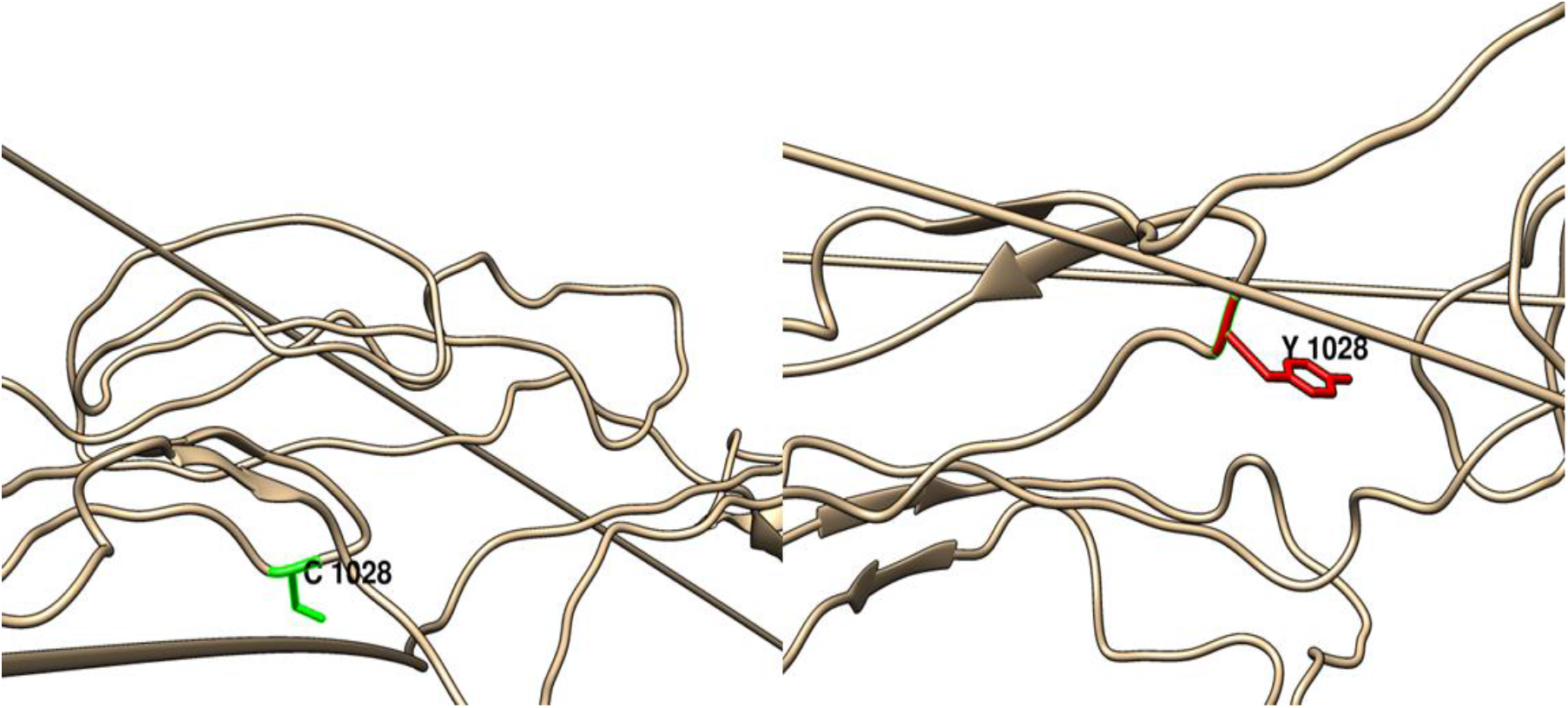
(C1028Y): change in the amino acid Cysteine (green box) into Tyrosine (red box) at position 1028.

**Figure (48):**
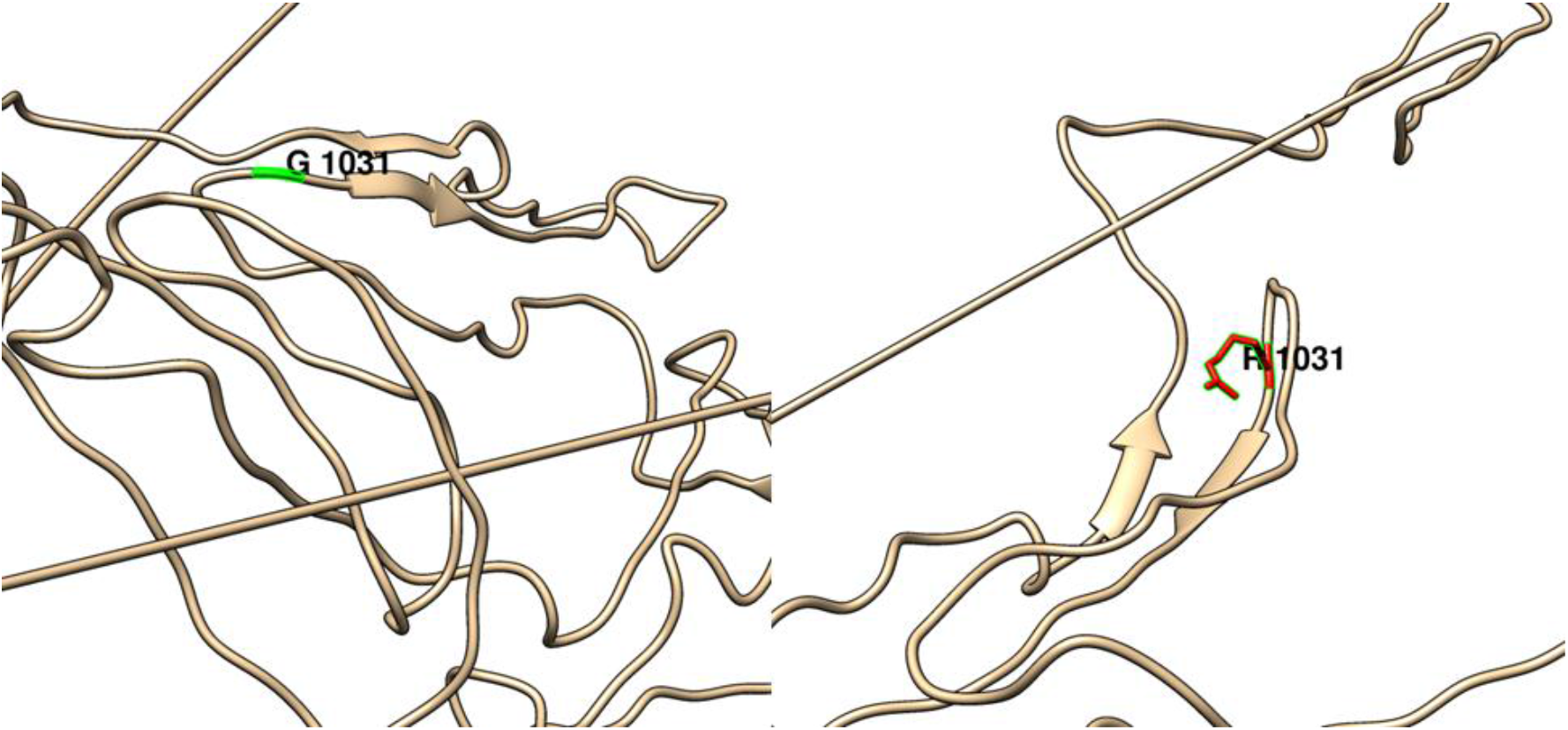
(G1031R): change in the amino acid Glycine (green box) into Arginine (red box) at position 1031.

**Figure (49):**
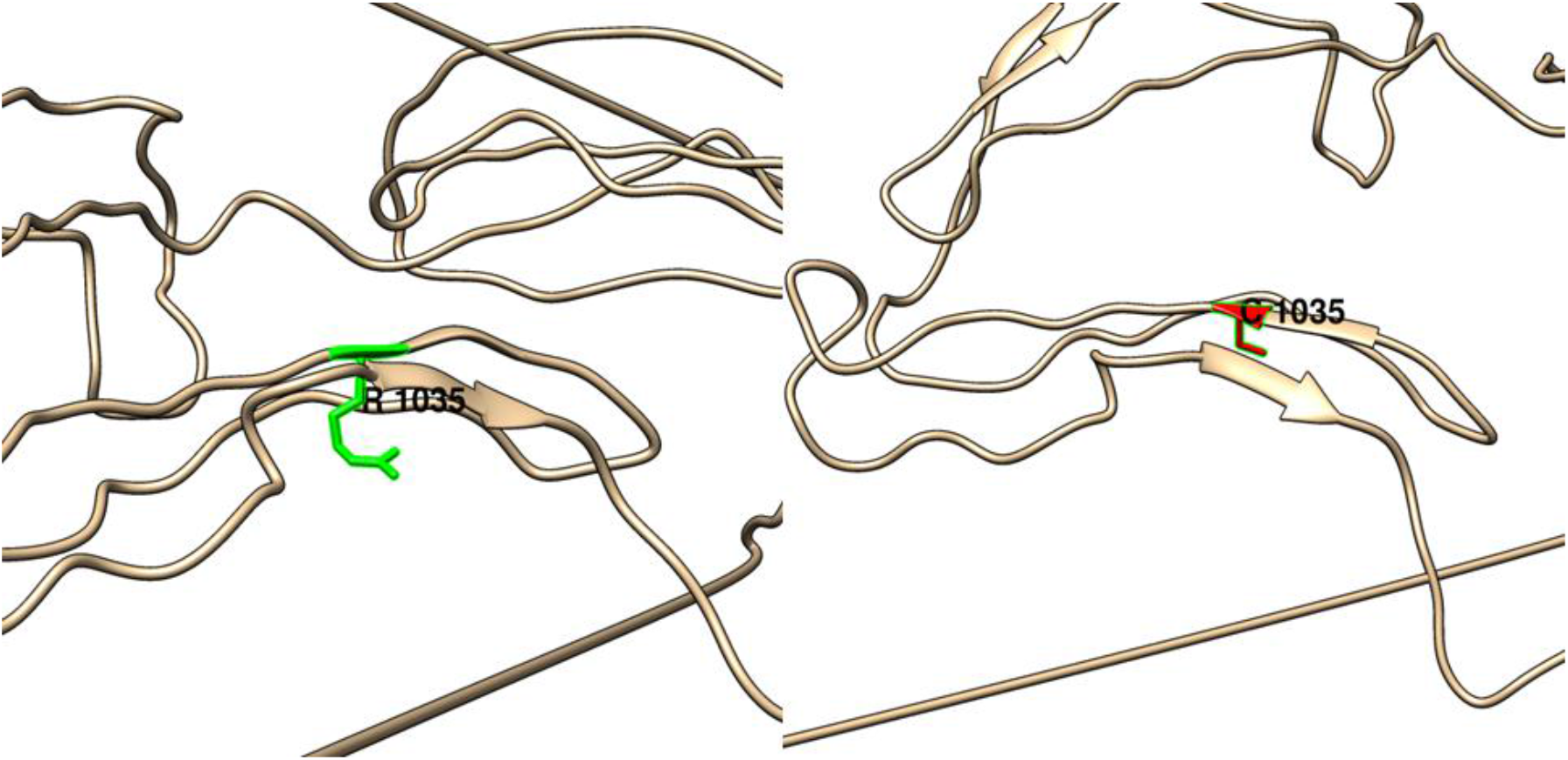
(R1035C): change in the amino acid Arginine (green box) into Cysteine (red box) at position 1035.

**Figure (50):**
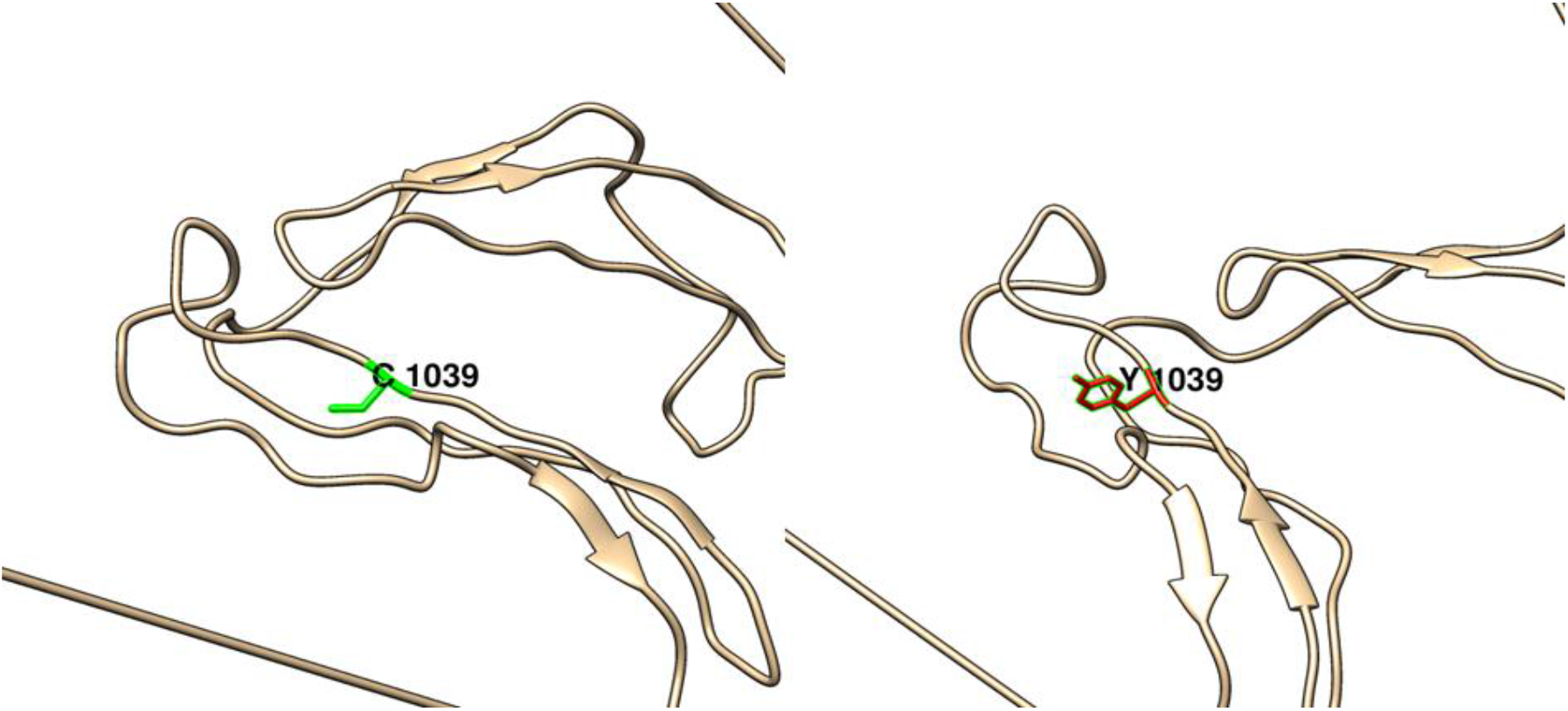
(C1039Y): change in the amino acid Cysteine (green box) into Tyrosine (red box) at position 1039.

**Figure (51):**
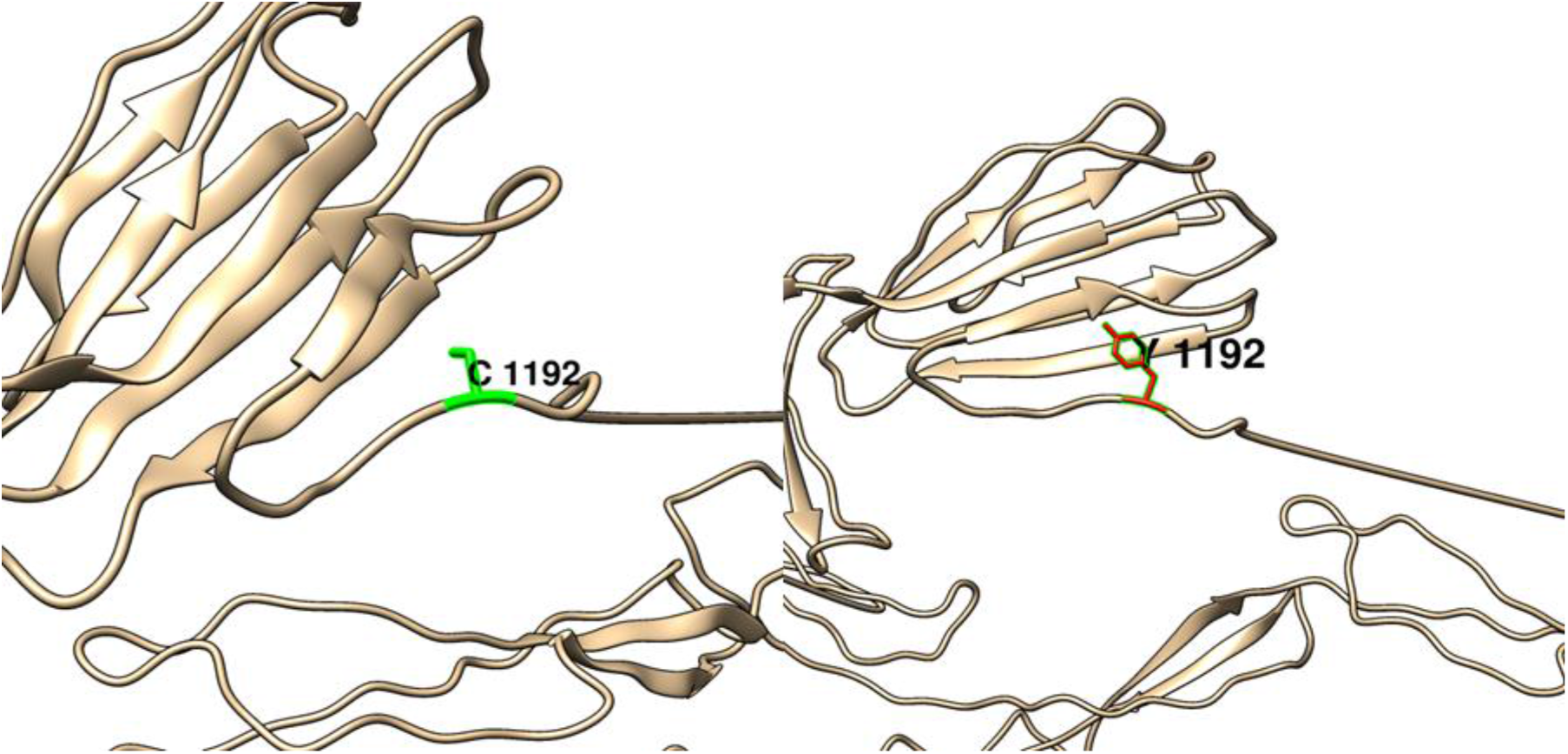
(C1192Y): change in the amino acid Cysteine (green box) into Tyrosine (red box) at position 1192.

### Interactions of *ADAMTS13* gene with other Functional Genes illustrated by GeneMANIA

GeneMANIA revealed that *ADAMTS13* has this function: metallopeptidase activity. The genes co-expressed with, share similar protein domain, or participate to achieve similar function are illustrated by GeneMANIA and shown in figure (52) Table (4) and (5).

**Table (4):** The *ADAMTS13* gene functions and its appearance in network and genome:

| Function | FDR | Genes in network | Genes in genome |
| --- | --- | --- | --- |
| metal ion transmembrane transporter activity | 0.813275 | 4 | 238 |
| inorganic cation transmembrane transporter activity | 0.813275 | 4 | 266 |
| metallopeptidase activity | 0.813275 | 3 | 80 |
| blood coagulation, intrinsic pathway | 0.813275 | 2 | 18 |
| blood coagulation, fibrin clot formation | 0.890003 | 2 | 21 |

\***FDR:** false discovery rate is greater than or equal to the probability that this is a false positive.

**Table (5):** The gene co-expressed, share domain and Interaction with *ADAMTS13* gene network

| Gene 1 | Gene 2 | Weight | Network group | Network |
| --- | --- | --- | --- | --- |
| KCNC4 | F8 | 0.008013 | Co-expression | Wang-Maris-2006 |
| ANGPTL6 | ADAMTS13 | 0.011534 | Co-expression | Mallon-McKay-2013 |
| TRPV1 | ADAMTS13 | 0.016478 | Co-expression | Mallon-McKay-2013 |
| POFUT2 | ADAMTS13 | 0.024919 | Co-expression | Mallon-McKay-2013 |
| C2orf72 | ADAMTS13 | 0.024335 | Co-expression | Mallon-McKay-2013 |
| KCNC4 | ADAMTS13 | 0.021195 | Co-expression | Mallon-McKay-2013 |
| MSI1 | ADAMTS13 | 0.021491 | Co-expression | Mallon-McKay-2013 |
| MSI1 | CACNG4 | 0.016442 | Co-expression | Mallon-McKay-2013 |
| KCNK17 | ANGPTL6 | 0.011594 | Co-expression | Mallon-McKay-2013 |
| CACNG4 | ADAMTS13 | 0.018641 | Co-expression | Dobbin-Giordano-2005 |
| GK2 | ADAMTS13 | 0.014198 | Co-expression | Dobbin-Giordano-2005 |
| TAS2R7 | ADAMTS13 | 0.018503 | Co-expression | Dobbin-Giordano-2005 |
| ADAMTS6 | ADAMTS13 | 0.010844 | Co-expression | Dobbin-Giordano-2005 |
| SLC6A13 | ADAMTS13 | 0.013106 | Co-expression | Dobbin-Giordano-2005 |
| AP4E1 | ADAMTS13 | 0.024775 | Co-expression | Dobbin-Giordano-2005 |
| AOC2 | GK2 | 0.018189 | Co-expression | Dobbin-Giordano-2005 |
| ANGPTL6 | ADAMTS13 | 0.030756 | Co-expression | Innocenti-Brown-2011 |
| TRPV1 | ADAMTS13 | 0.012766 | Co-expression | Innocenti-Brown-2011 |
| KCNK17 | ADAMTS13 | 0.025007 | Co-expression | Innocenti-Brown-2011 |
| POFUT2 | ADAMTS10 | 0.011832 | Co-expression | Noble-Diehl-2008 |
| AOC2 | ADAMTS13 | 0.014705 | Co-expression | Noble-Diehl-2008 |
| ADAMTS7 | ADAMTS13 | 0.025154 | Co-expression | Smirnov-Cheung-2009 |
| GK2 | ADAMTS13 | 0.019954 | Co-expression | Smirnov-Cheung-2009 |
| SLC6A9 | ADAMTS13 | 0.022157 | Co-expression | Smirnov-Cheung-2009 |
| SLC6A13 | ADAMTS13 | 0.016187 | Co-expression | Smirnov-Cheung-2009 |
| CLDN6 | ADAMTS13 | 0.020231 | Co-expression | Smirnov-Cheung-2009 |
| ADAMTS7 | ADAMTS13 | 0.011667 | Co-expression | Wang-Cheung-2015 |
| TAS2R7 | ADAMTS13 | 0.011175 | Co-expression | Wang-Cheung-2015 |
| SLC6A9 | ADAMTS13 | 0.012103 | Co-expression | Wang-Cheung-2015 |
| AOC2 | ADAMTS13 | 0.016991 | Co-expression | Wang-Cheung-2015 |
| CLDN6 | ADAMTS13 | 0.010212 | Co-expression | Wang-Cheung-2015 |
| CACNG4 | ADAMTS13 | 0.023497 | Co-expression | Chen-Brown-2002 |
| ADAMTS10 | ADAMTS13 | 0.022796 | Co-expression | Chen-Brown-2002 |
| AP4E1 | ADAMTS10 | 0.016491 | Co-expression | Chen-Brown-2002 |
| SLC6A13 | SLC6A9 | 0.003413 | Co-expression | Wu-Garvey-2007 |
| F8 | VWF | 0.060129 | Pathway | Wu-Stein-2010 |
| F8 | VWF | 0.294935 | Pathway | REACTOME |
| VWF | ADAMTS13 | 0.55051 | Physical Interactions | IREF-DIP |
| F8 | ADAMTS13 | 0.387079 | Physical Interactions | IREF-DIP |
| F8 | VWF | 0.387079 | Physical Interactions | IREF-DIP |
| F8 | VWF | 0.185599 | Physical Interactions | BIOGRID-SMALL-SCALE-STUDIES |
| VWF | ADAMTS13 | 0.72218 | Physical Interactions | IREF-INTACT |
| VWF | ADAMTS13 | 0.586387 | Physical Interactions | IREF-HPRD |
| F8 | VWF | 0.081584 | Physical Interactions | IREF-HPRD |
| VWF | ADAMTS13 | 0.692532 | Predicted | Wu-Stein-2010 |
| ADAMTS7 | ADAMTS13 | 0.01789 | Shared protein domains | INTERPRO |
| ADAMTS10 | ADAMTS13 | 0.017896 | Shared protein domains | INTERPRO |
| ADAMTS10 | ADAMTS7 | 0.0318 | Shared protein domains | INTERPRO |
| ADAMTS6 | ADAMTS13 | 0.017888 | Shared protein domains | INTERPRO |
| ADAMTS6 | ADAMTS7 | 0.031785 | Shared protein domains | INTERPRO |
| ADAMTS6 | ADAMTS10 | 0.031796 | Shared protein domains | INTERPRO |
| SLC6A13 | SLC6A9 | 0.054687 | Shared protein domains | INTERPRO |
| CLDN6 | CACNG4 | 0.015305 | Shared protein domains | INTERPRO |
| ADAMTS7 | ADAMTS13 | 0.020768 | Shared protein domains | PFAM |
| ADAMTS10 | ADAMTS13 | 0.018679 | Shared protein domains | PFAM |
| ADAMTS10 | ADAMTS7 | 0.025162 | Shared protein domains | PFAM |
| ADAMTS6 | ADAMTS13 | 0.018676 | Shared protein domains | PFAM |
| ADAMTS6 | ADAMTS7 | 0.025158 | Shared protein domains | PFAM |
| ADAMTS6 | ADAMTS10 | 0.030488 | Shared protein domains | PFAM |
| SLC6A13 | SLC6A9 | 0.052632 | Shared protein domains | PFAM |
| CLDN6 | CACNG4 | 0.022244 | Shared protein domains | PFAM |

**Figure (52):**
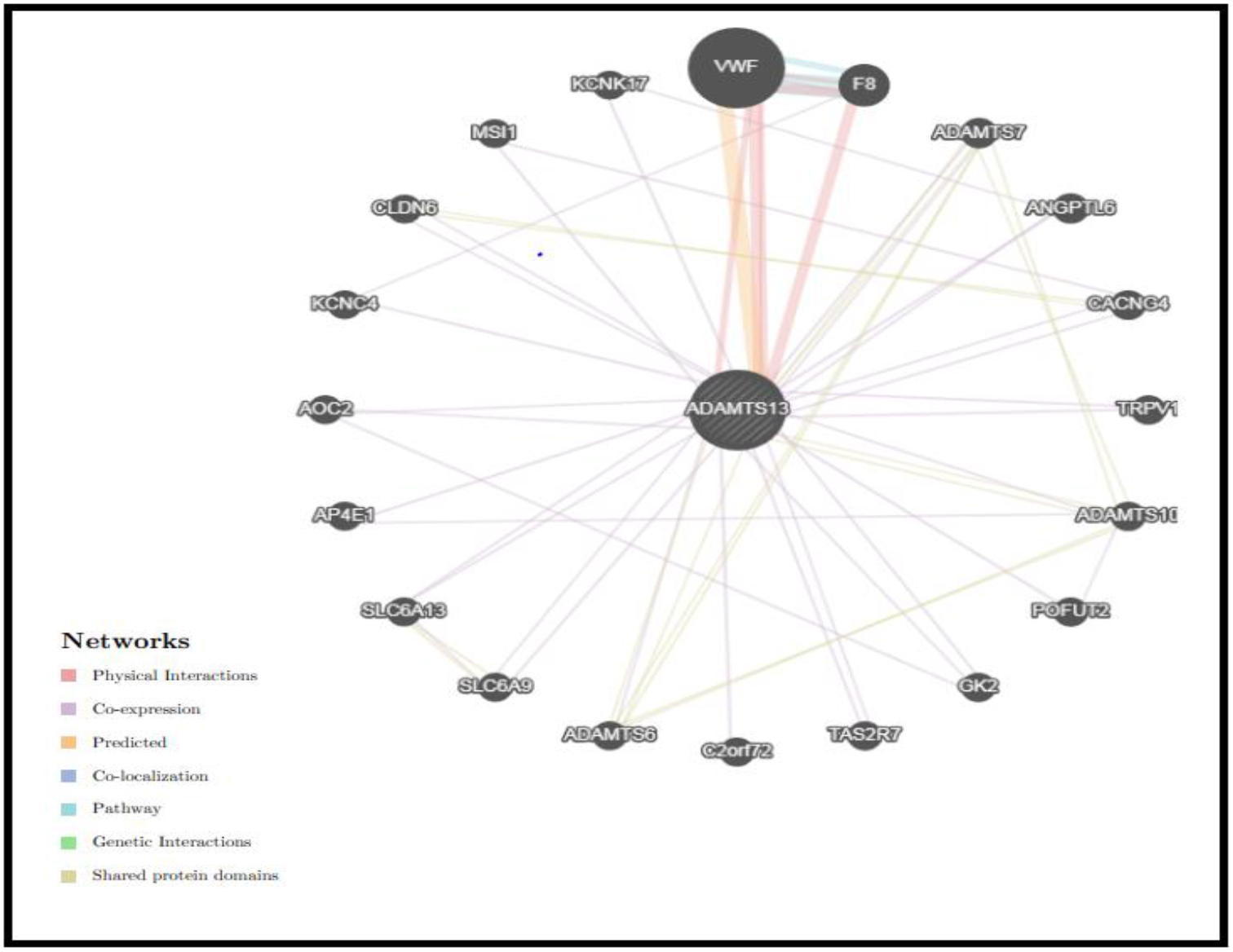
Interaction between *ADAMTS13* and its related genes

### Novel deleterious SNPs in *ADAMTS13* gene revealed by this study

**Table (6):**
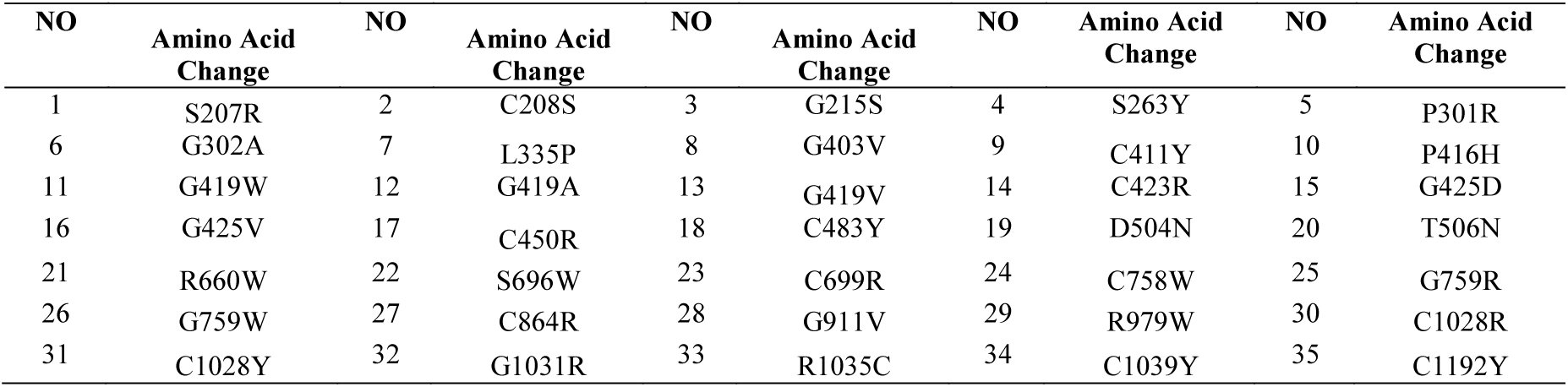
Novel deleterious mutations in *ADAMTS13* gene

| NO | Amino Acid Change | NO | Amino Acid Change | NO | Amino Acid Change | NO | Amino Acid Change | NO | Amino Acid Change |
| --- | --- | --- | --- | --- | --- | --- | --- | --- | --- |
| 1 | S207R | 2 | C208S | 3 | G215S | 4 | S263Y | 5 | P301R |
| 6 | G302A | 7 | L335P | 8 | G403V | 9 | C411Y | 10 | P416H |
| 11 | G419W | 12 | G419A | 13 | G419V | 14 | C423R | 15 | G425D |
| 16 | G425V | 17 | C450R | 18 | C483Y | 19 | D504N | 20 | T506N |
| 21 | R660W | 22 | S696W | 23 | C699R | 24 | C758W | 25 | G759R |
| 26 | G759W | 27 | C864R | 28 | G911V | 29 | R979W | 30 | C1028R |
| 31 | C1028Y | 32 | G1031R | 33 | R1035C | 34 | C1039Y | 35 | C1192Y |

## DISCUSSION

Our in-depth analysis of the SNPs in the *ADAMTS13* gene found 51 amino acid changes that are highly pathogenic according to our insilico analysis. According to the dbSNP database some of these SNPs are listed as per their clinical impact as benign, pathogenic, and untested or no information is available. One of the aims of our study is to confirm or refute these claims based on computational analysis. 35 deleterious SNPs, not previously mentioned in the literature, were determined to be novel mutations and are listed in Table (6).

R398H, C508Y, C951G, C1024G are listed as being pathogenic by dbSNP. Several studies [14, 21, 22, 36–38] have found R398H mutation to be detrimental. Similarly, wet lab studies confirmed the pathogenicity of C508Y[21], C951G and C1024G[36] mutations. H234Q, D235Y, C311Y, P353L, W390C, G525D and C908Y are categorized as ‘untested’ in dbSNP. H234Q mutation in the *ADAMTS13* gene found to be damaging through insilico methods in this study was also found to the cause of repeated renal failure in an adult patient [39]. D235Y mutation has also been studied in patients with congenital TTP [40]. Similarly, C311Y and P353L mutations in the *ADAMTS13* gene have been found in patients with congenital TTP in more than one study[14, 37]. W390C mutation has been implicated in TTP and HUM (hemolytic uremic syndrome) [5]. G525D and C908Y *ADAMTS13* mutations were found in Japanese patients with Upshaw-Shulman syndrome [2]. No clinical information was provided by dbSNP for the following mutations: R398C, R409W, R421C, G761S and C908S. However, several studies have indeed been conducted. A paper published in 2012 studied congenital TTP patients in the UK and confirmed the pathogenicity of R398C and R409W mutations and their impact on *ADAMTS13* activity [22]. Sequencing of the *ADAMTS* gene in patients with deep vein thrombosis identified R421C mutation [41]. G761S mutation was identified in patients with impaired renal function [40], while C908S mutation was one of 10 *ADAMTS13* gene mutations in six families with congenital TTP [38].

## CONCLUSION

According to our analysis, we found 35 nsSNPs effects on *ADAMTS13* protein leading to thrombotic thrombocytopenic purpura using computational approach. Bioinformatics tools are vital in prediction analysis, making use of increasingly voluminous biomedical data thereby providing markers for screening or for genetic mapping studies.

## ACKNOWLEDGMENT

The authors wish to acknowledgment the enthusiastic cooperation of Africa City of Technology – Sudan.

## CONFLICT OF INTEREST

The authors declare that there is no conflict of interest regarding the publication of this paper.

## DATA AVAILABILITY

All relevant data used to support the findings of this study are included within the manuscript and supplementary information files.

## Notes

https://www.ncbi.nlm.nih.gov/

